# Historic museum samples provide evidence for a recent replacement of *Wolbachia* types in European *Drosophila melanogaster*

**DOI:** 10.1101/2023.06.25.546460

**Authors:** Anton Strunov, Sandra Kirchner, Julia Schindelar, Luise Kruckenhauser, Elisabeth Haring, Martin Kapun

## Abstract

*Wolbachia* is one of the most common bacterial endosymbionts, which is frequently found in numerous arthropods and nematode taxa. *Wolbachia* infections can have a strong influence on the evolutionary dynamics of their hosts since these bacteria are reproductive manipulators that affect the fitness and life history of their host species for their own benefit. Host-symbiont interactions with *Wolbachia* are perhaps best studied in the model organism *Drosophila melanogaster*, which is naturally infected with five different types among which wMel and wMelCS are the most frequent ones. Comparisons of infection types between natural flies and long-term lab stocks have previously indicated that wMelCS represents the ancestral type, which was only very recently replaced by the nowadays dominant wMel in most natural populations. In this study, we took advantage of recently sequenced museum specimens of *D. melanogaster* that have been collected 90-200 years ago in Northern Europe to test this hypothesis. Our comparison to contemporary *Wolbachia* samples provides compelling support for the replacement hypothesis and identifies potential infections with yet unknown *Wolbachia* types of supergroup B. Our analyses show that sequencing data from historic museum specimens and their bycatch are an emerging and unprecedented resource to address fundamental questions about evolutionary dynamics in host-symbiont interactions.

## 1. Introduction

*Wolbachia pipientis* (Hertig, 1936) is a gram-negative ⍺-proteobacterium of the order Rickettsiales, which represents one of the most common endosymbionts in animals (reviewed in Landmann, 2019; Kaur *et al*., 2021). *Wolbachia* has been detected in approximately 50% of all arthropods and some filarial nematode species (Zug & Hammerstein, 2012; Weinert *et al*., 2015; Lefoulon *et al*., 2016; Sazama *et al*., 2017) and can have a substantial impact on the life history and fitness of its host. These effects might be transient (Correa & Ballard, 2016) and range from parasitic to mutualistic with different fitness effects on the host (Zug & Hammerstein, 2015) depending on the host species, the *Wolbachia* type and the environment (reviewed in Pietri *et al*., 2016; Chrostek *et al*., 2021; Strunov *et al*., 2022). Moreover, *Wolbachia* can act as a reproductive parasite causing cytoplasmic incompatibility, feminization, male-killing and parthenogenesis (reviewed in Landmann, 2019; Kaur *et al*., 2021). Since *Wolbachia* is maternally transmitted, these reproductive manipulations usually lead to a higher prevalence and increased fitness of infected females in a population (reviewed in Werren *et al*., 2008). Moreover, by enhancing host fitness and fecundity (Dedeine *et al*., 2001; Hosokawa *et al*., 2010; Miller *et al*., 2010) and by providing protection against RNA viruses (Hedges *et al*., 2008; Teixeira *et al*., 2008; Moreira *et al*., 2009; Osborne *et al*., 2009) Wolbachia can behave as facultative or obligate mutualist (Zug & Hammerstein, 2015). *Wolbachia* thus plays an important role in the evolution of its host species (reviewed in Kaur *et al*., 2021) and can even lead to speciation (Miller *et al*., 2010).

Currently, *Wolbachia* is classified into several phylogenetic supergroups traditionally subsumed under *W. pipientis* (Bandi *et al*., 1994; Werren *et al*., 1995; Lo *et al*., 2002; Olanratmanee *et al*., 2021). However, systematics of the genus *Wolbachia* is still under debate and there is no consensus yet of whether supergroups represent distinct species or just lineages of *W. pipientis* (e.g., Ramírez-Puebla *et al*., 2015; Lindsey *et al*., 2016 and references therein). In the following we thus use *Wolbachia* synonymously for *W. pipientis*. Most *Wolbachia* infecting insects belong to supergroups A and B (Lo *et al*., 2002; Gerth *et al*., 2014). However, due to horizontal transfers across species boundaries, the phylogenetic tree of *Wolbachia* is often incongruent with the species tree of their hosts (Gomes *et al*., 2022). In fact, several insect host species, such as the vinegar fly *Drosophila simulans* can even carry *Wolbachia* variants that belong to different supergroups, such as type wRi (Hoffmann *et al*., 1986) and wHa (O’Neill & Karr, 1990) from supergroup A and type wNo from supergroup B (Poinsot & Merçot, 1997). However, with the exception of *D. mauritania,* whose *Wolbachia* symbiont is of supergroup B (Meany *et al*., 2019), most other species of the genus *Drosophila* have *Wolbachia* types that belong to supergroup A only. The genetic model species *Drosophila melanogaster*, for example, has been found to carry at least five distinct *Wolbachia* types of supergroup A in natural populations (Riegler *et al*., 2012). A large body of literature documents that specifically the two most common natural types wMel and wMelCS in *D. melalnogaster*, but also wMelPop, which evolved under laboratory conditions from a wMelCS background (Min & Benzer, 1997; Woolfit *et al*., 2013), can have very distinct fitness effects on their hosts (e.g., Serga *et al*., 2021; Strunov *et al*., 2022). For example, wMelCS is usually characterized by higher bacterial titers (Chrostek *et al*., 2013) and can influence reproduction and other life history traits depending on the environment and the host genetic background (Strunov *et al*., 2022).

The wMel type is nowadays the most common natural type in worldwide *D. melanogaster* populations, whereas wMelCS only occurs in a few populations at very low frequencies (Richardson *et al*., 2012; Chrostek *et al*., 2013). However, comparisons of *Wolbachia* types in recent populations and long-standing *D. melanogaster* labstrains revealed that old labstocks, which were often collected almost a century ago, were all dominated by the wMelCS type (Riegler *et al*., 2005). These findings indicate that wMel only recently replaced the ancestral wMelCS type in worldwide host populations, within the last 50 to 100 years (Riegler *et al*., 2005; Nunes *et al*., 2008; Richardson *et al*., 2012; Early & Clark, 2013; Ilinsky, 2013). Potential biological reasons for such a rapid turn-over are still under debate. Higher titers in wMelCS (Chrostek *et al*., 2013) could result in putatively higher fitness costs to the host, e.g., due to reduced fecundity (see, for example, Serga *et al*., 2014), which may have facilitated the spread of the low-titer wMel type. Alternatively, threats by RNA viruses may have changed over time, which could have reduced the benefit of higher virus protection by wMelCS (Hedges *et al*., 2008; Teixeira *et al*., 2008; Moreira *et al*., 2009; Osborne *et al*., 2009; Chrostek *et al*., 2021) and subsequently benefitted the spread of wMel.

Furthermore, given that no data of historic samples from this time period were available until now, it remains unclear to which extent the findings in Riegler et al. (2005) reflect the true evolutionary history of both types in natural populations. For example, labstocks may be prone to contamination in stock centers, which could have biased their infection type if wMelCS is more successful than wMel under laboratory conditions. This could have accordingly resulted in an excess of wMelCS infections in labstrains and thus bias the interpretation of these data. Moreover, the origin and pervasiveness of the invasion and subsequent replacement are also under debate since residual occurrence of wMelCS infections in natural populations have been identified in Europe and North America (Richardson *et al*., 2012).

In our study, we are for the first time able to empirically address this long-standing question by taking advantage of a recently published genomic dataset from *Drosophila melanogaster* museum specimens, which were collected 90-200 years ago in Northern Europe (Shpak *et al*., 2023). We tested for the presence of *Wolbachia*-specific reads in these data, estimated titer variation and investigated the relatedness to contemporary *Wolbachia* strains. This unique dataset, which we complemented with contemporary genomic data from various sources, showed, that *D. melanogaster* populations in Northern Europe were indeed dominated by the ancestral wMelCS types up to 90 years ago which further supports the hypothesis that wMel only recently replaced wMelCS in worldwide populations.

## 2. Materials and Methods

In total, 67 genomic datasets were included in the present study (Table 1): Recently published whole genomic Illumina deep sequencing data of 25 historic *D. melanogaster* museum specimens (Shpak *et al*., 2023) were tested for the presence of *Wolbachia*-specific reads. These samples originated from Sweden, Denmark and Germany and had been collected between 90 and 200 years ago. Complementary to these historic samples, we used Oxford Nanopore sequencing technology (ONT) to newly sequence genomic DNA of six freshly collected isofemale lines from wild populations in Portugal and Finland, that were naturally infected with either the wMel or the wMelCS *Wolbachia* types and three labstocks that had been artificially infected with wMel, wMelCS and wMelPOP (Truitt *et al*., 2019; see Table 1). Furthermore, we included 24 raw Illumina sequencing datasets of contemporary *D. melanogaster* specimens, which were previously tested positive for infections with wMel or wMelCS (Richardson *et al*., 2012), for phylogenetic analyses (see Table 1). These samples are part of the *Drosophila* NEXUS datasets (Pool *et al*., 2012; Lack *et al*., 2015, 2016) and the DGRP datasets (Mackay *et al*., 2012), which represent huge collections of single individual sequencing data of natural flies that were mostly collected in Africa and North America, respectively. Since these datasets contained only two specimens infected with wMelCS, we further included the raw sequencing datasets of two labstocks carrying wMelCS (Woolfit *et al*., 2013; Duarte *et al*., 2021). Complementary to these raw sequencing data, we further obtained nine RefSeq assemblies (Wu *et al*., 2004; Singhal & Mohanty, 2018; Basting & Bergman, 2019; Duarte *et al*., 2021) for phylogenetic inference based on BUSCO genes - eight of them carrying wMel and one them a wMelCS type (see Table 1). Given the close relationship between wYak (the *Wolbachia* strain of *Drosophila yakuba*) and wMel (Scholz *et al*., 2020), we used the RefSeq sequence information of the wYak *Wolbachia* type (NZ_VCEF01000001.1) as an outgroup for phylogenetic tree reconstructions.

**Table 1.**
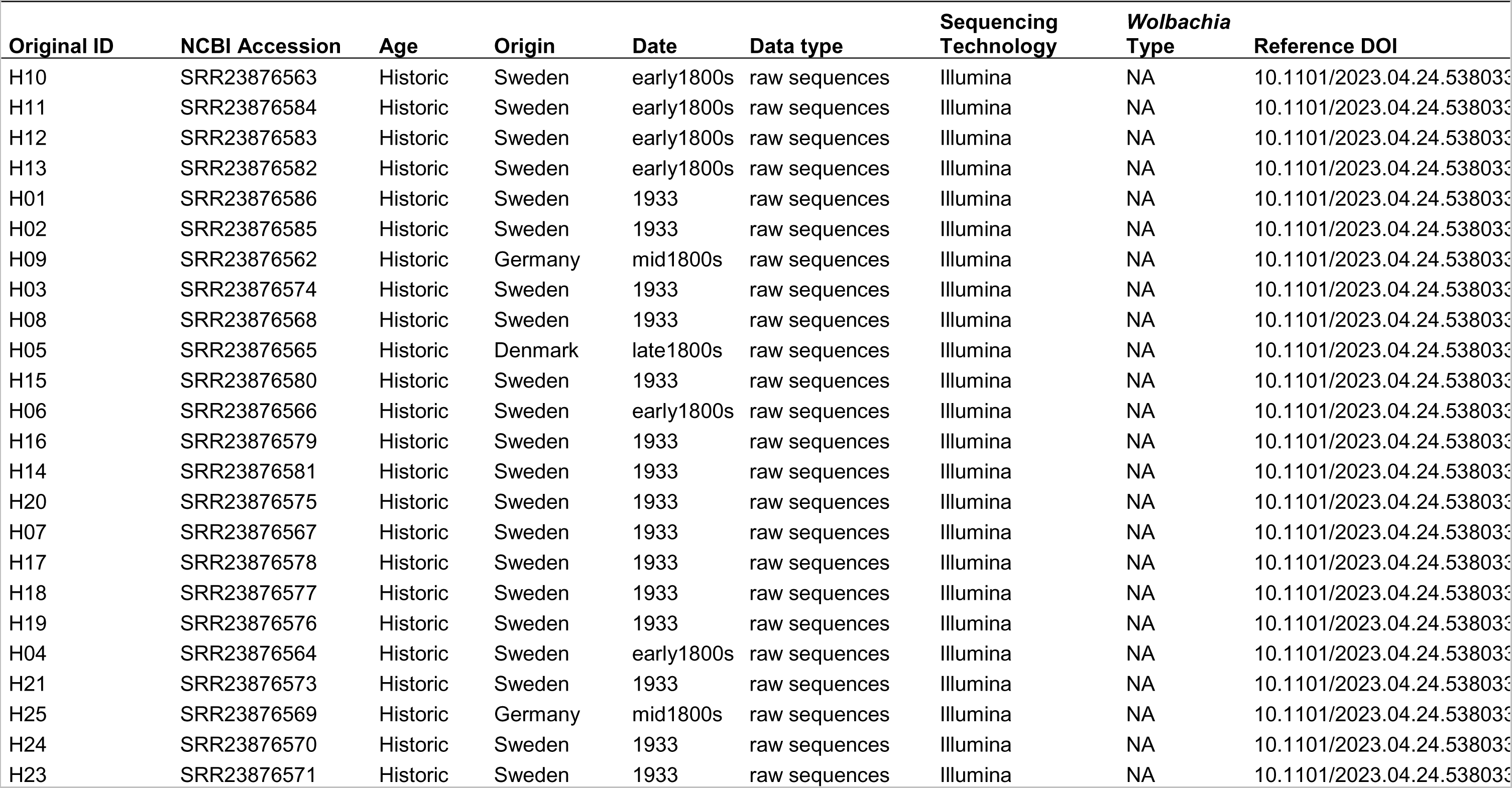
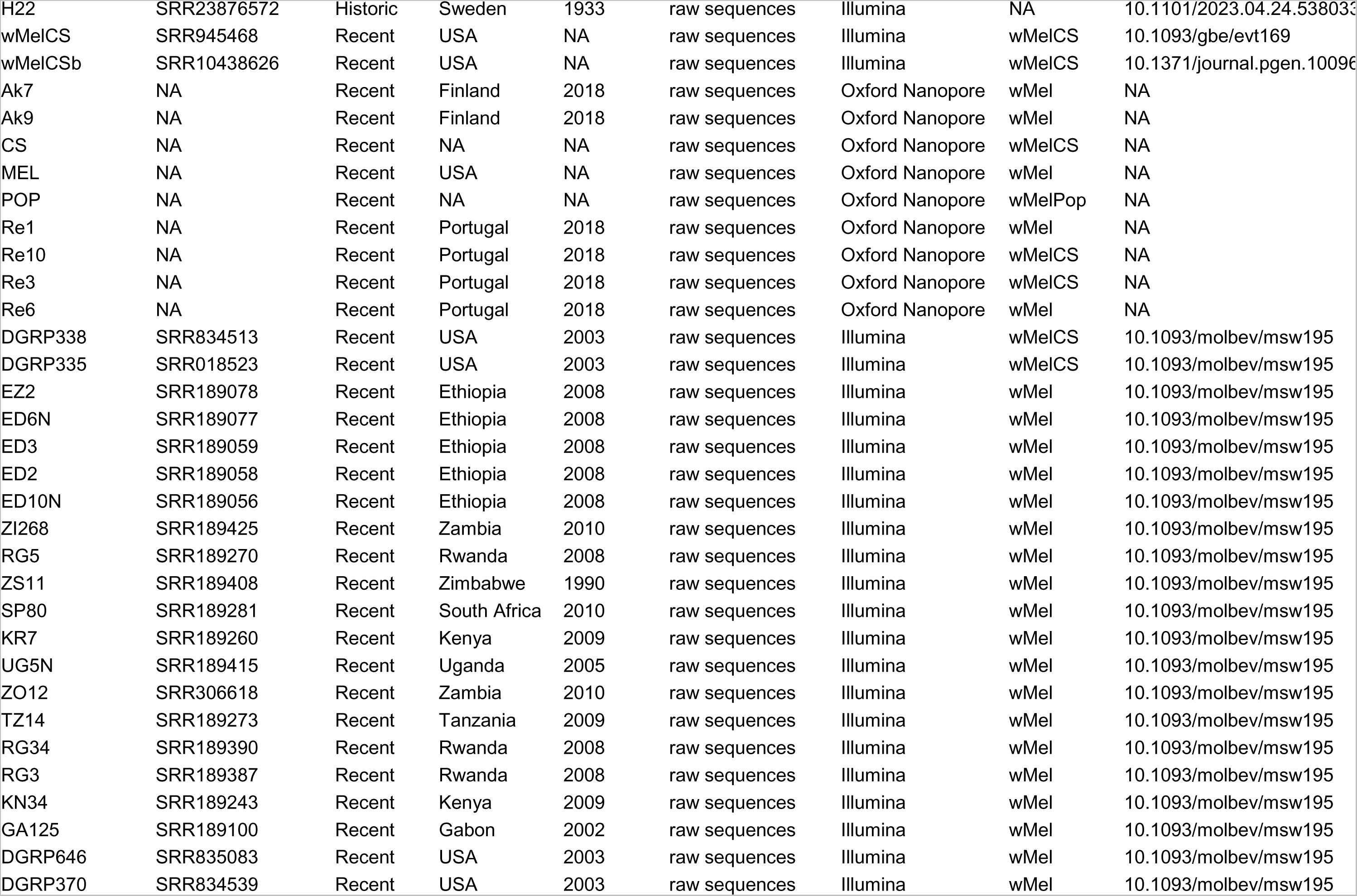
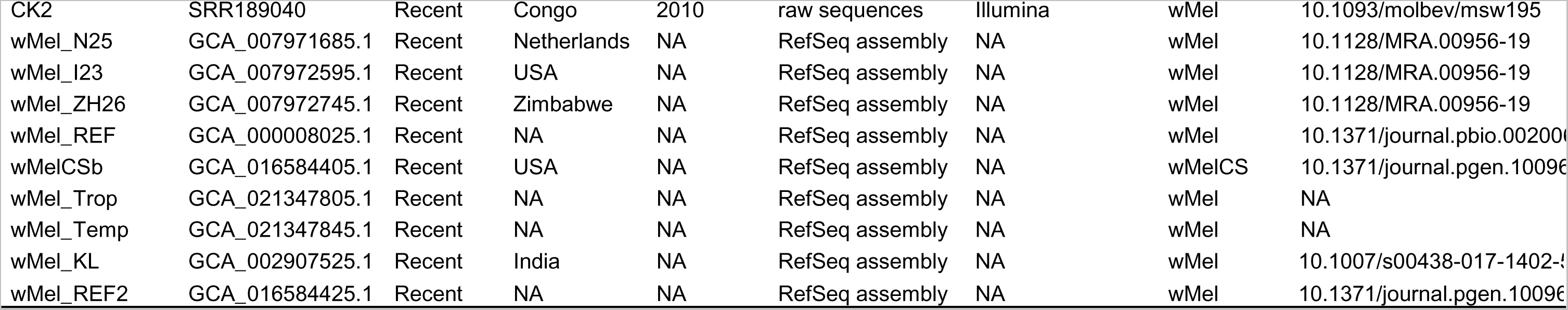
Summary table of sequencing data used in this study. Whenever available, this table is showing sample-specific information on sampling location and collection year as well as technical information such as data type, sequencing technology, infection type, NCBI accession number and DOI of the associated studies. Note, the upper part of the table contains raw sequencing data of different sources, while the last part includes nine assembled Wolbachia genomes of the RefSeq database.

### 2.1 DNA extraction, library preparation and ONT whole genome sequencing

Prior to whole genome sequencing of nine contemporary isofemale lines (Table 1) infected with either wMel, wMelCS or wMelPOP by means of ONT technology, we confirmed their infection status and infection type with PCR using *Wolbachia* type-specific VNTR-141 primers and PCR conditions as described in Riegler et al. (2012). Then, we isolated high molecular weight DNA by following the Monarch T3010 DNA purification kit (NEB, USA) protocol. For each of the highly inbred strains, we pooled 25 females prior to DNA extraction to increase the DNA yield, assuming that the amount of residual heterozygosity in the strains is very low. After DNA extraction, the DNA was stored at 8°C for 7 days in order to ensure relaxation and homogenization of the extracts. Prior to library preparations, the integrity/purity of the extracted DNA (A260/A280 and A260/A230 ratios) was assessed with a BioPhotometer (Eppendorf) and quantification of the DNA yield was measured via Qubit Fluorometer (ThermoFisher Scientific; dsDNA Broad Range Assay Kit). The fragment length distribution of the extracted DNA was inspected on a TapeStation system (Agilent) and extracted DNA was subjected to library preparation by applying the Native Barcoding Kit 24 Sequencing Kit (SQK-NBD112.24) for genomic DNA from Oxford Nanopore Technologies. For library preparation we strictly followed the manufacturer’s protocol using 400 ng of gDNA per sample as input. The protocol included a repair and end-prep step, followed by barcode ligation, and subsequent adapter ligation and clean-up step of the pooled samples. After finishing the library preparation, app. 11 fmol of DNA library were loaded on a Flow Cell (R9, FLO-MIN106D) and sequencing was performed on a MinION Mk1C (Oxford Nanopore Technologies) for 72h. Since the yield of *Wolbachia*-specific reads of all wMel-type samples was markedly lower than of wMelCS-type libraries, we repeated ONT sequencing of the five wMel-type samples multiplexed on a new flow cell. We then performed base calling and demultiplexing of the samples using the GPU version of guppy (v. 6.2.1; Wick *et al*., 2019).

### 2.2 Characterization of Wolbachia infections: relative titer and infection status

Prior to all downstream analyses we trimmed the raw Illumina reads of historic and contemporary samples using trimgalore (v 0.6.2; Martin, 2011) allowing a minimum base quality of 20. We only retained intact read pairs with a minimum length of 30bp for further analyses. To obtain estimates of *Wolbachia* titers in historic and contemporary samples, we used bwa mem (v. 0.7.13; Li, 2013) for Illumina sequencing data or minimap2 (v. 2.17; Li, 2018) for ONT sequencing data to map all raw FASTQ reads from each sample against a joint reference sequence, which was constructed from the *Drosophila melanogaster* reference genome v.6 (Hoskins *et al*., 2015) and additional genome sequences of other common microbial symbionts, including the wMel reference genome (see Kapun *et al*., 2020 for more details). Using the command *samtools coverage* of the samtools program (v. 1.12; Li *et al*., 2009) in combination with a custom *python* script (SumReadDepths.py), we calculated average read depths (RD) for all *Drosophila* chromosomes and symbiont genomes. Based on this information, we estimated relative *Wolbachia* titers for a given sample by dividing the average RD at the *Wolbachia* genome by the average RD across all *Drosophila* autosomes (see also Scholz *et al*., 2020). Based on average RD and the proportion of the *Wolbachia* genome covered by reads, we classified *Drosophila* samples into three infection statuses: (1) infected with *Wolbachia* (>50% of the *Wolbachia* genome covered by reads; >10-fold average RD; relative titer >0.3:1), (2) uninfected (<15% of the *Wolbachia* reference genome covered by reads; <2-fold average RD; relative titer < 0.01:1), or (3) with unclear status (15-30% of the *Wolbachia* reference genome covered by reads; 2 to 9-fold average RD; relative titer 0.02:1 to 0.2:1).

### 2.3. Classification based on diagnostic SNPs and BLAST searches

We identified 67 SNPs in 33 contemporary samples that were fixed for different alleles in the 25 wMel and the eight wMelCS samples, respectively (Table 1). Then, we inferred the allelic state for these diagnostic SNPs in 13 of the historic samples that were putatively infected with *Wolbachia.* We only considered a position whenever the RD was ≥ 2 and the allelic state was unambiguously identified based on the posterior probability as explained above. Three of the samples with uncertain infection status (H02, H06 and H22) did not have coverage at any of the diagnostic markers and thus had to be excluded.

Since an unambiguous *Wolbachia* classification failed for samples H03 and H05, we pre-filtered with Kraken (v. 2.1.2; Wood & Salzberg, 2014) using a custom-built database that consisted of the published genomes of wMel (RefSeq: AE017196.1), wMelCS (RefSeq: NZ_JACSNK000000000.1) and wMelPOP (RefSeq: NZ_AQQE00000000.1) and only retained reads which matched the references in the database. Then, we blasted 100,000 randomly drawn raw forward reads against a local copy of the NCBI nt database using the *blastn* algorithm (v.2.12.0; Camacho *et al*., 2009). We only considered the best match per read passing stringent similarity thresholds (i.e., *e*-value < 1e-50; >99% sequence similarity) and specifically focused on matches with *Wolbachia*-specific target sequences. Based on the target sequence ID (sseqid) and the NCBI TaxID (staxids), we manually obtained information on the corresponding supergroup for each of the matched *Wolbachia* types from current literature (Werren *et al*., 1995; Lo *et al*., 2002; Wu *et al*., 2004; Saha *et al*., 2012; Scholz *et al*., 2020; Olanratmanee *et al*., 2021; Neupane *et al*., 2022). In addition, we obtained the reference genome of wMeg, a *Wolbachia* type of the bowfly *Chrysomya megacephala* from supergroup B (RefSeq: NZ_CP021120.1) which yielded most BLAST hits for both samples, and mapped raw reads against a joint reference consisting of the genomes of wMeg and wMelCS to test if the reads of H03 and H05 preferentially map to the supergroup B reference genome. Furthermore, we included five additional samples that were unambiguously identified as wMelCS as controls in our analysis.

### 2.4 De-Novo assembly of historic and contemporary Wolbachia genomes

In a next step, we used raw FASTQ files of each library that were trimmed and prefiltered with Kraken as described above for de-novo assembly with SPAdes (v. 3.15.3; Bankevich *et al*., 2012; Prjibelski *et al*., 2020) using default parameters. We further used the combined raw long-fragment reads from two rounds of ONT sequencing for contemporary flies infected with wMel or the reads of a single round of ONT sequencing from wMelCS- or wMelPop-infected flies for assembly with Flye (v. 2.9; Lin *et al*., 2016; Kolmogorov *et al*., 2019) using default parameters. Raw assemblies were finally polished in two rounds using Racon (v. 1.5.0; Vaser *et al*., 2017)

Subsequently, assembly quality was assessed based on common quality statistics, such as numbers of contigs, N50 and N90 with QUAST (v. 5.1.0rc1; Mikheenko *et al*., 2018). We tested for assembly completeness using the BUSCO approach (v. 5.2.2; Seppey *et al*., 2019; Manni *et al*., 2021), where the proportion of intact, fragmented, and missing benchmarking universal single-copy orthologs (BUSCO) specific to the bacterial order Rickettsiales (rickettsiales_odb10) was evaluated in each assembled genome. In addition, we re-mapped the raw reads to the assembled contigs using minimap2 (v. 2.17; Li, 2018) to assess variation in read-depth and we compared all contigs to a local copy of the NCBI *nt* database using *blastn* of the BLAST suite (v.2.12.0; Camacho *et al*., 2009). After that, we visualized the results of these quality assessments with Blobtools (v. 3.0.0; Laetsch & Blaxter, 2017).

Finally, we used the published *Wolbachia* wMel reference genome (RefSeq: AE017196.1) as a backbone to align and orient the raw contigs with *nucmer* of the MUMmer package (v. 3.23; Marçais *et al*., 2018). Based on *show-tiling* of the MUMmer package, we identified the minimum number of unique contigs that span a maximum of the reference backbone. Using two custom python scripts (CombineContigs.py and SetStart.py), we then combined these contigs into a single scaffold and filled the gaps between each pair of consecutive contigs with a string of ten N’s. Moreover, given that the bacterial genome is circular, we anchored the newly assembled scaffolds at the start point of the reference genome and shifted protruding sequences to the end of the scaffolds.

### 2.5 Phylogenetic analysis

We employed two complementary approaches to explore the evolutionary history of *Wolbachia* based on phylogenetic inference.

#### 2.5.1 SNP-based phylogenetic analysis

First, we mapped raw Illumina reads of *Wolbachia*, which were pre-filtered with Kraken, as explained above, against the wMel reference genome (RefSeq: AE017196.1). For the Illumina sequencing data of historic and contemporary samples downloaded from NCBI SRA, we mapped paired-end reads using bwa mem (v. 0.7.13; Li, 2013) with default parameters. Conversely, we used minimap2 (v. 2.17; Li, 2018) with default parameters to map long-fragment reads from ONT sequencing against the reference *Wolbachia* genome. Raw BAM files were filtered with samtools (v. 1.12; Li *et al*., 2009) to contain mapped reads only (parameter -F 4) and sorted by reference position using the *samtools sort* command.

Then, we used the BCFtools (v. 1.16; Danecek *et al*., 2021) command *bcftools mpileup* to synchronize the mapped reads of all samples and called SNPs using *bcftools call* assuming haploidy and stored the types in the VCF file format. Using a custom Python script (BCF2Phylip.py) we converted the VCF file to the PHYLIP format, only considering bi-allelic polymorphic positions where the posterior probability of the most likely genotype was > 50 and of the alternative genotype < 30. In addition, we only included positions with more than 5-fold RD in each of the samples and removed samples, with more than 50% missing information across all SNPs. We used a total of 279 aligned SNP data to reconstruct a maximum likelihood (ML) tree based on the GTR-CAT substitution model from 20 randomly drawn maximum parsimony (MP) starting trees using RaXML (v. 2.8.10; Kozlov *et al*., 2019) and additionally performed 100 rounds of bootstrapping to test for the robustness of nodes. The final tree was plotted in *R* (v. 4.2.2; R Core Team, 2019) using the *ggtree* package (Yu *et al*., 2017).

#### 2.5.2 Phylogenetic analysis based on BUSCO genes

Complementary to the approach described above, we compared the nucleotide sequences of candidate genes as identified by the BUSCO approach from the de novo assembled genomes of the historic museum specimens and the contemporary samples. We therefore focused on 104 BUSCO genes specific to the bacterial order Rickettsiales, which were identified as complete and which were present in the majority of the assembled genomes in our dataset. Using MAFFT with standard parameters (v. 7.487; Katoh & Standley, 2013), we aligned their sequences separately and subsequently concatenated the alignments with a custom Python script (ConcatenateAlignments.py; final length of concatenated alignment: 68,992bp) Similar to above, we reconstructed a ML tree based on the GTR-Gamma substitution model from 20 distinct randomized MP starting trees and additionally performed 100 rounds of bootstrapping to test for the robustness of nodes using RaXML (v. 2.8.10; Kozlov *et al*., 2019). The final tree was plotted in *R* (v. 4.2.2; R Core Team, 2019) using the *ggtree* package (Yu *et al*., 2017).

### 2.6 Comparison to mitochondrial phylogeny

*Wolbachia* and the host mitochondria are both transmitted maternally to the offspring and should thus share a similar evolutionary history (Nunes *et al*., 2008; Richardson *et al*., 2012; Ilinsky, 2013). This allows testing for genomic signals of horizontal introgression of *Wolbachia* into a host, which would manifest in inconsistencies among the species trees of mitochondria and *Wolbachia*. We therefore employed the SNP-based phylogenetic approach described above also based on mitochondrial reads, which we pre-filtered by comparing all raw reads against a custom-built Kraken database consisting of the *D. melanogaster* reference mitochondrial genome (RefSeq: NC_024511.2). To this end, we included all historic samples irrespective of their infection status. We then performed a phylogenetic analysis with this data set using RaXML as explained above.

In addition, we used a subset of the *Wolbachia* and mitochondrial SNP datasets that included the same samples and performed a phylogenetic analysis as described above. Then, We used the function *as.dendrogram* from the *R* package *phylogram* (Wilkinson & Davy, 2018) to convert the *Wolbachia* and mitochondrial tree files in NEWICK format to ultrametric trees. We first untangled the two trees, i.e., we swapped the branches to best fit the order of the samples using the *untangle* function with the “step1side” method of the *dendextend* package (Galili, 2015) and then produced a tanglegram using the *tanglegram* function to visualize the relationship between the two trees.

## 3. Results & Discussion

In this study we took advantage of a recently published genomic dataset consisting of 25 museum samples of *D. melanogaster* collected between 200 and 90 years ago in Northern Europe. Besides testing if museomics of century-old *Drosophila* samples allows to identify historic *Wolbachia* infections, we address several long-standing questions concerning the co-evolution of *D. melanogaster* and *Wolbachia*. In particular, these data for the first time allow to test the hypothesis that the historically dominating wMelCS *Wolbachia* type, nowadays found world-wide only in a few populations at low frequencies, was only recently replaced by the wMel type within the last century or whether the wMel type was already present in European populations centuries ago.

### 3.1 Wolbachia infections in several historic Drosophila samples

As a first step, we classified the 25 sequenced historic *Drosophila* samples as infected or uninfected based on reference mapping. Similar to Shpak et al. (2023), we found that the average read length of the historic samples after trimming was approximately 50 bp (Table 2). However, we observed that in spite of our rigorous quality filtering, RD at the *Drosophila* genome were markedly higher in our study compared to the results in Shpak et al. (2023). We assume that this discrepancy is the result of differences in the mapping pipelines that we employed. Specifically, in contrast to Shpak et al. (2023), we used the more recent *D. melanogaster* reference genome v.6 and the *bwa mem* mapping algorithm. A rigorous visual comparison of mapped reads of historic and contemporary samples with IGViewer (v. 2.10.2; Robinson *et al*., 2011) did not reveal any obvious mapping errors. However, we noted that forward and reverse reads were perfectly overlapping in the historic samples, which is expected given the short expected fragment size of app. 50bp (Shpak *et al*., 2023). Given that we only considered intact read pairs, we do not believe that this introduces a bias in our analyses.

**Table 2.** Mapping statistics of raw Illumina reads from historic and contemporary samples. Table showing properties of the samples, i.e., their sampling age, their infection status and infection type as well as average read lengths after trimming, read depths (RD) at *Drosophila* autosomes, X chromosome, mitochondrial genome and *Wolbachia* reference genomes. In addition, the last two columns provide the proportion of the *Wolbachia* genome that is covered by reads and the relative *Wolbachia* titer, which is the ratio of average RD from *Wolbachia* and *Drosophila* autosomes.

| Sample ID | Age | *Wolbachia* Type | Infection Status | Average Read length [bp] | Read Depth [fold] |  |  |  | *Wolbachia* reference coverage [%] | *Wolbachia* relative titer |
| --- | --- | --- | --- | --- | --- | --- | --- | --- | --- | --- |
|  |  |  |  |  | Autosomes | X | Mitochondrion | *Wolbachia* |  |  |
| H03 | Historic | unclear | unclear | 56.9 | 41.9 | 23.2 | 4798.8 | 8.0 | 45.9 | 0.190 |
| H05 | Historic | unclear | unclear | 68.1 | 53.6 | 37.8 | 2527.7 | 4.9 | 37.5 | 0.092 |
| H02 | Historic | wMelCS | unclear | 66.4 | 3.9 | 4.0 | 410.9 | 0.2 | 1.7 | 0.040 |
| H06 | Historic | wMelCS | unclear | 54.4 | 23.8 | 23.1 | 4181.8 | 2.3 | 13.0 | 0.098 |
| H10 | Historic | wMelCS | unclear | 47.5 | 133.7 | 75.8 | 675.5 | 3.8 | 32.6 | 0.029 |
| H13 | Historic | wMelCS | unclear | 46.6 | 97.5 | 54.8 | 600.1 | 1.7 | 16.6 | 0.017 |
| H22 | Historic | wMelCS | unclear | 58.8 | 41.4 | 41.9 | 2893.3 | 2.3 | 34.4 | 0.055 |
| H04 | Historic | wMelCS | infected | 53.0 | 39.6 | 39.8 | 3288.0 | 20.4 | 59.6 | 0.515 |
| H07 | Historic | wMelCS | infected | 48.6 | 97.1 | 101.9 | 8872.6 | 208.5 | 99.8 | 2.149 |
| H09 | Historic | wMelCS | infected | 47.1 | 186.3 | 197.2 | 85591.3 | 223.7 | 99.9 | 1.201 |
| H11 | Historic | wMelCS | infected | 45.8 | 144.3 | 81.6 | 1036.0 | 47.1 | 95.1 | 0.327 |
| H12 | Historic | wMelCS | infected | 45.7 | 91.9 | 93.0 | 1960.4 | 54.3 | 88.7 | 0.590 |
| H15 | Historic | wMelCS | infected | 48.6 | 126.0 | 71.0 | 12320.3 | 737.8 | 100.0 | 5.856 |
| H16 | Historic | wMelCS | infected | 46.5 | 229.1 | 137.8 | 11955.7 | 1018.0 | 100.0 | 4.443 |
| H20 | Historic | wMelCS | infected | 46.1 | 109.5 | 114.8 | 8431.8 | 316.5 | 100.0 | 2.890 |
| H23 | Historic | wMelCS | infected | 64.0 | 86.7 | 90.5 | 5656.8 | 498.3 | 100.0 | 5.746 |
| H24 | Historic | wMelCS | infected | 57.0 | 50.3 | 51.9 | 5940.3 | 136.6 | 100.0 | 2.716 |
| H01 | Historic |  | uninfected | 55.1 | 52.3 | 29.1 | 2797.9 | 0.4 | 5.0 | 0.008 |
| H08 | Historic |  | uninfected | 59.0 | 72.1 | 40.8 | 4324.7 | 1.3 | 10.4 | 0.018 |
| H14 | Historic |  | uninfected | 46.4 | 195.6 | 110.9 | 12841.6 | 0.3 | 2.8 | 0.001 |
| H17 | Historic |  | uninfected | 47.1 | 124.2 | 72.1 | 8967.1 | 0.2 | 3.2 | 0.002 |
| H18 | Historic |  | uninfected | 62.6 | 66.4 | 66.7 | 3400.5 | 0.3 | 3.4 | 0.005 |
| H19 | Historic |  | uninfected | 47.9 | 208.3 | 119.5 | 11642.9 | 0.5 | 6.1 | 0.002 |
| H21 | Historic |  | uninfected | 54.3 | 24.5 | 13.6 | 1091.0 | 0.0 | 1.1 | 0.001 |
| H25 | Historic |  | uninfected | 53.6 | 48.2 | 51.9 | 4540.6 | 0.2 | 6.3 | 0.005 |
| Ak7_full | Recent | wMel | infected | NA | 3.9 | 4.1 | 210.1 | 17.2 | 100.0 | 4.434 |
| Ak9_full | Recent | wMel | infected | NA | 4.0 | 4.3 | 383.7 | 25.7 | 100.0 | 6.400 |
| MEL_full | Recent | wMel | infected | NA | 3.1 | 3.2 | 245.7 | 14.2 | 100.0 | 4.648 |
| Re1_full | Recent | wMel | infected | NA | 2.9 | 2.9 | 211.7 | 13.6 | 100.0 | 4.738 |
| Re6_full | Recent | wMel | infected | NA | 4.7 | 5.0 | 395.9 | 20.1 | 100.0 | 4.315 |
| CK2 | Recent | wMel | infected | 69.0 | 40.5 | 38.4 | 562.3 | 38.2 | 99.9 | 0.942 |
| ED10N | Recent | wMel | infected | 36.8 | 34.9 | 34.4 | 132.4 | 12.0 | 99.6 | 0.344 |
| ED2 | Recent | wMel | infected | 36.4 | 27.5 | 28.6 | 19.4 | 5.9 | 97.3 | 0.213 |
| ED3 | Recent | wMel | infected | 36.5 | 26.9 | 25.5 | 237.6 | 13.8 | 99.8 | 0.514 |
| ED6N | Recent | wMel | infected | 36.9 | 33.1 | 34.2 | 50.6 | 30.5 | 100.0 | 0.920 |
| EZ2 | Recent | wMel | infected | 36.6 | 28.0 | 26.7 | 34.0 | 34.8 | 100.0 | 1.239 |
| GA125 | Recent | wMel | infected | 37.4 | 39.5 | 38.9 | 99.5 | 25.4 | 99.9 | 0.643 |
| KN34 | Recent | wMel | infected | 36.5 | 26.1 | 24.1 | 561.6 | 27.8 | 99.9 | 1.065 |
| KR7 | Recent | wMel | infected | 36.6 | 25.4 | 24.5 | 169.1 | 15.8 | 99.8 | 0.621 |
| RG3 | Recent | wMel | infected | 37.0 | 34.8 | 32.3 | 360.5 | 30.0 | 100.0 | 0.863 |
| RG34 | Recent | wMel | infected | 37.1 | 27.5 | 28.5 | 110.6 | 10.3 | 99.7 | 0.375 |
| RG5 | Recent | wMel | infected | 36.3 | 22.6 | 24.3 | 71.0 | 568.2 | 100.0 | 25.192 |
| SP80 | Recent | wMel | infected | 67.8 | 35.9 | 33.0 | 68.6 | 15.7 | 99.7 | 0.436 |
| TZ14 | Recent | wMel | infected | 36.9 | 32.7 | 33.5 | 85.3 | 58.2 | 100.0 | 1.782 |
| UG5N | Recent | wMel | infected | 36.9 | 30.4 | 31.4 | 49.8 | 47.3 | 100.0 | 1.554 |
| ZI268 | Recent | wMel | infected | 37.4 | 38.9 | 37.1 | 346.8 | 67.7 | 100.0 | 1.739 |
| ZO12 | Recent | wMel | infected | 49.0 | 44.2 | 42.1 | 57.3 | 38.8 | 100.0 | 0.877 |
| ZS11 | Recent | wMel | infected | 37.0 | 33.1 | 34.8 | 16.0 | 70.2 | 100.0 | 2.123 |
| CS | Recent | wMelCS | infected | NA | 1.3 | 1.4 | 99.1 | 15.2 | 99.9 | 11.711 |
| DGRP335 | Recent | wMelCS | infected | 21.5 | 2.4 | 1.9 | 56.9 | 21.0 | 100.0 | 8.909 |
| DGRP338 | Recent | wMelCS | infected | 33.0 | 11.7 | 9.5 | 45.5 | 65.9 | 100.0 | 5.611 |
| Re10 | Recent | wMelCS | infected | NA | 1.6 | 1.7 | 163.0 | 14.6 | 100.0 | 9.084 |
| Re3 | Recent | wMelCS | infected | NA | 1.2 | 1.2 | 130.0 | 19.2 | 100.0 | 15.433 |
| POP | Recent | wMelPop | infected | NA | 1.1 | 1.2 | 116.9 | 49.4 | 100.0 | 43.087 |

We found that absolute *Wolbachia*-specific read-depths (RD) varied between basically zero to thousand-fold across samples (see Table 2). Similarly, we observed large variations in the length of the reference sequence that was covered by reads. Proportions were ranging from as low as 0.9% to 100%. Since absolute *Wolbachia*-specific RDs are strongly influenced by overall sample-specific RD, we further calculated relative *Wolbachia* titers by dividing *Wolbachia*-specific RD by average *Drosophila*-specific RDs at the four autosomal arms (2L, 2R, 3L and 3R). *Wolbachia* titers relative to the *Drosophila* host in museum samples varied dramatically and ranged from 0.001:1 up to 5:1 (Figure 1).

**Figure 1.**
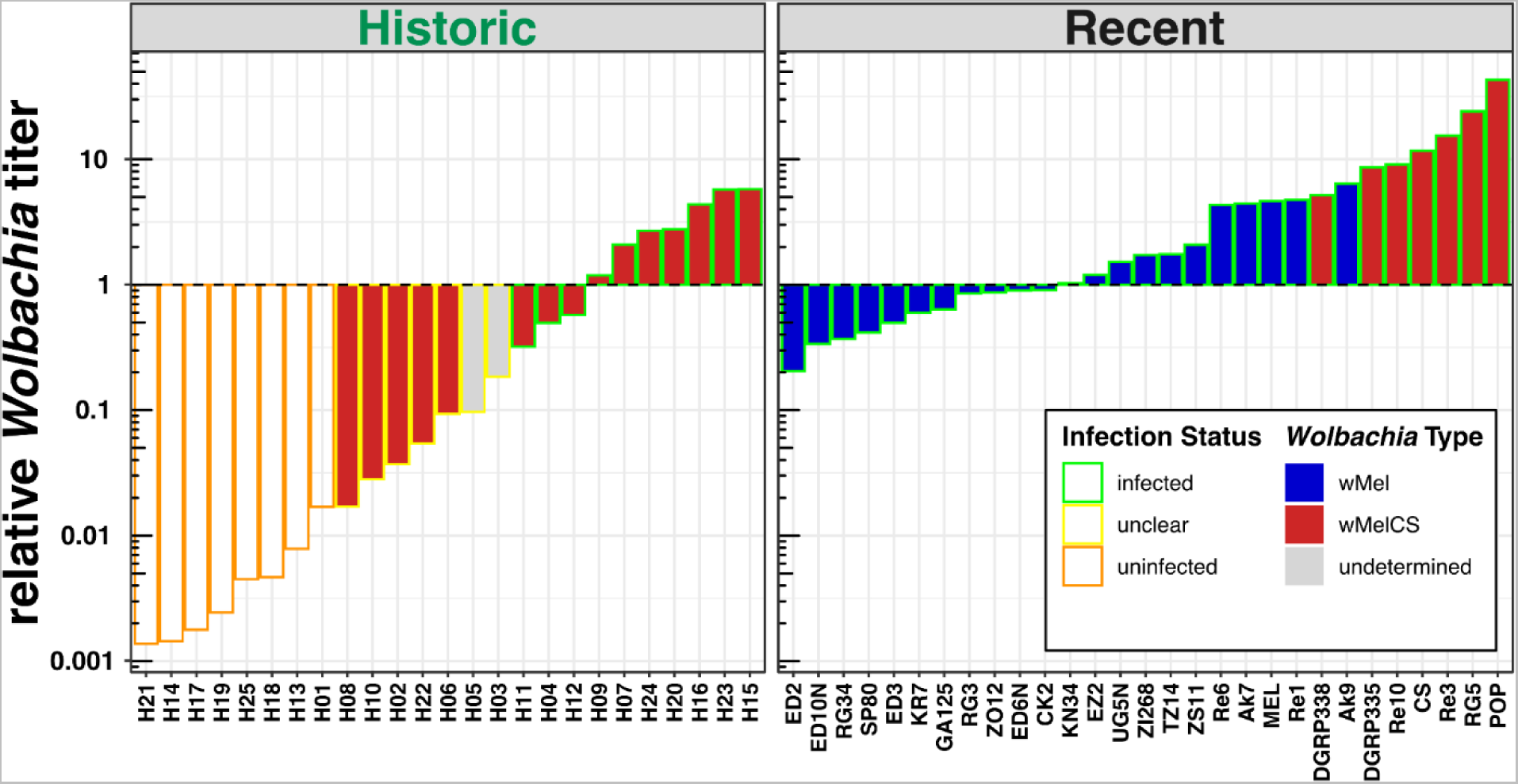
Relative *Wolbachia* titers. Barplots showing the ratio of *Wolbachia*-specific reads to *Drosophila*-specific reads for each library in historic and contemporary samples. The color outline of each bar indicates the inferred infection status of historic and contemporary samples, where infected specimens are shown in green, uninfected specimens are highlighted in orange and flies with uncertain infection status are shown in yellow. The fill color depicts the *Wolbachia* type in the recent samples. Here and in all other figures, wMel is shown in blue and both the wMelCS and the wMelPOP type, are shown in red. The yet undetermined *Wolbachia* type in the putatively infected historic samples H03 and H05 is shown in gray.

Based on these results, we qualitatively classified 10 historic samples as unambiguously infected with *Wolbachia* (see Material & Methods for evaluation criteria). Conversely, we considered eight historic samples as uninfected. Besides the two aforementioned infection types, we identified seven samples with uncertain infection status, which were characterized by low RD (2 to 9-fold), which covered only parts (15-30%) of the reference genome and which had quite low relative titers (0.02:1 to 0.2:1).

Surprisingly, all samples that we classified as uninfected contained even small proportions of reads that mapped to the *Wolbachia* genome. We thus cannot rule out that the low bacterial titers and RDs in putatively uninfected samples or samples with uncertain infection status are in fact false negatives. Low titers may have a biological explanation if *Wolbachia* titers in natural samples have been lower 100-200 years ago. Alternatively, the age of the specimen may play a role if in museum samples *Wolbachia*-specific DNA decays at different rates than *Drosophila*-specific DNA. DNA shearing, depurination and deamination are major sources of degradation in air-dried museum specimens (Zimmermann *et al*., 2008; Raxworthy & Smith, 2021). With age, the highest decay is observed in dG content (Zimmermann *et al*., 2008), which might explain unequal degradation of bacterial and eukaryotic DNA depending on GC content. Low titers may also represent an artifact of the DNA extraction or library preparation process, when uninfected flies or samples with uncertain infection status may have inadvertently been cross-contaminated from samples with very high bacterial titers that were processed at the same time (Raxworthy & Smith, 2021).

In contrast, we observed no difference in average relative titer in infected flies from historic and recent samples (paired *t*-test, *p*=0.07). In the contemporary samples, however, we found significant differences in relative titer between samples infected with wMel and wMelCS (paired *t-*test, *p*=0.0009). This finding is in good agreement with previous data from qPCR-based analyses of bacterial titers, which indicate that relative titers of wMelCS are markedly higher and in the order of ≤ 1:1 for *w*Mel and 1:1 to 10:1 for *w*MelCS types, depending on the age of the flies under laboratory conditions (Chrostek *et al*., 2013)

### 3.2 Wolbachia infections in historic D. melanogaster samples are mostly of type wMelCS and putatively of a yet unknown member of supergroup B

To test if the *Wolbachia* variants identified in the historic samples are of wMel and/or wMelCS type, we took advantage of our genomic dataset from recent samples with known *Wolbachia* type and identified 67 diagnostic marker SNPs that were fixed for different allelic states in the two types. We were able to investigate thirteen historic samples, where at least three SNPs contained allelic information due to sufficient RD. Eleven historic samples, which we found to be putatively infected with *Wolbachia*, were unambiguously of wMelCS type (Figure 2). Conversely, we found that samples H03 and H05 were completely ambiguous since approximately half of the covered SNPs with allelic information classified the samples as wMel and the other half as wMelCS (Figure 2). Diagnostic SNPs with wMel or wMelCS-specific alleles were randomly distributed along the *Wolbachia* reference genome, which strongly suggests that the observed heterogeneous allelic patterns are not the result of a recombination event among a wMel and a wMelCS type, which would have resulted in a contiguous stretch of type-specific diagnostic alleles.

**Figure 2.**
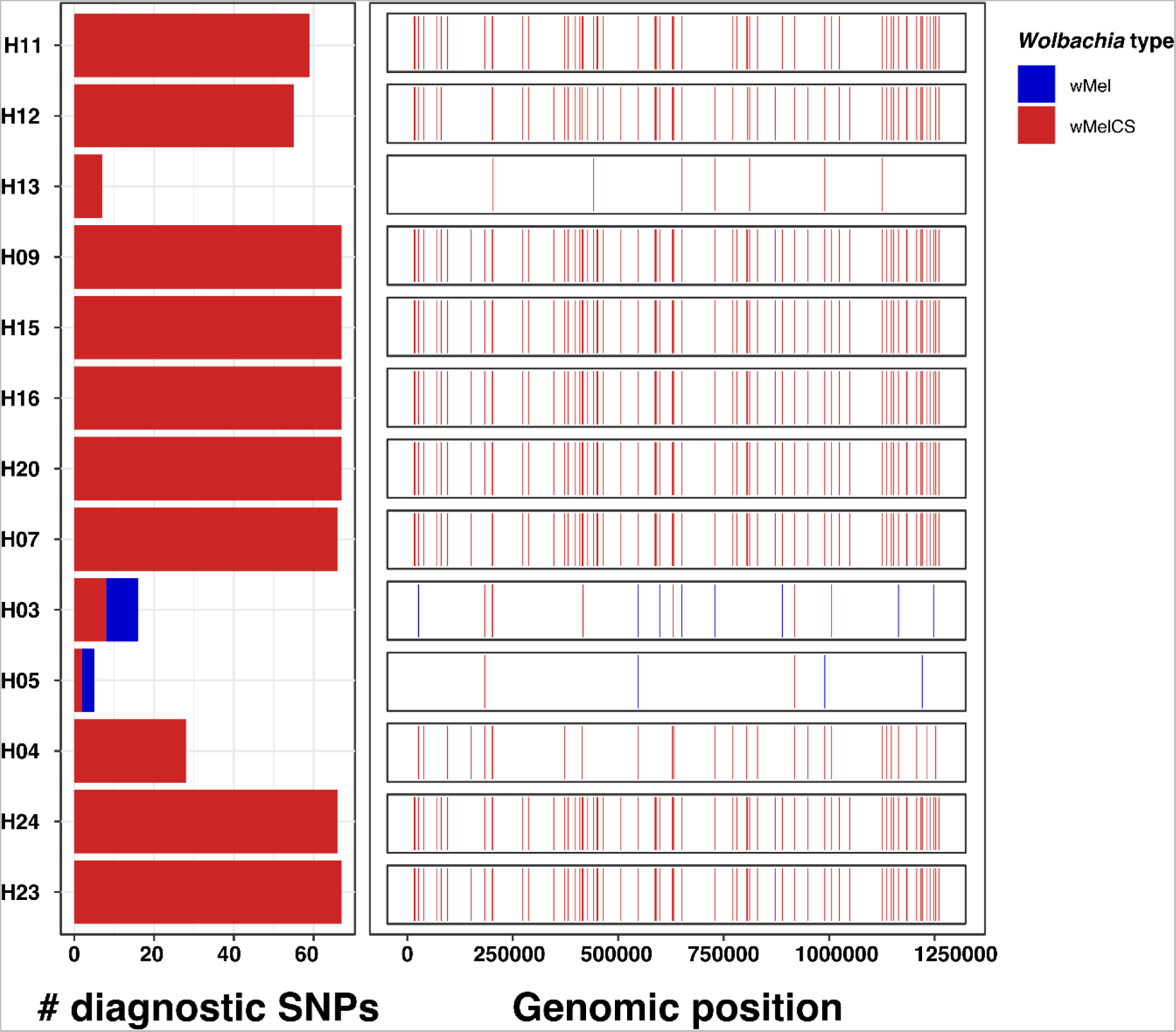
*Wolbachia* type characterization based on diagnostic marker SNPs. The left barplot shows the allelic states (blue: wMel, red: wMelCS) for a maximum of 67 diagnostic *Wolbachia*-type specific marker SNPs. The size of the bars indicates the number of SNPs with available allelic information for a given sample. The right panel shows the genomic distribution and allelic state of the diagnostic SNPs for a given sample. This figure includes 13 historic samples where allelic information was available for at least one diagnostic SNP.

However, while *Wolbachia* double-infection has not been described yet in *D. melanogaster*, but only in its sister species *D. simulans* (Rousset & Solignac, 1995), we cannot rule out that the observed pattern may indicate the presence of both types in sample H03. Given the fact that we assumed haploidy of *Wolbachia* in their hosts when we identified SNP variants, we would, however, fail to identify heterozygosities. We therefore repeated SNP-calling for these two samples, now also allowing for biallelic SNPs. As a control, we likewise called SNPs for sample H13, which had similarly low RD but was unambiguously characterized as wMelCS and thus showed no sign of double infections. Our analyses revealed that the histograms of allele frequency in putative heterozygous SNPs did not differ between the three datasets (paired *t*-test; *p* > 0.05 for all comparisons) which indicates that both uncertain samples were not co-infected with two *Wolbachia* types.

We further tested if the *Wolbachia* reads in the samples H03 and H05 were potentially more similar to yet another *Wolbachia* type by blasting raw reads against the nt database only taking into consideration hits with >99% sequence identity and e-values > 1e-50. While many BLAST hits for both samples matched *Wolbachia* (H03: 921 matches, 51.9% and H05: 80 matches,13%; Figure 3), we found that only very small proportions of *Wolbachia*-specific hits were matching wMel or wMelCS (H03: 15 matches; 1.63% and H05: 1 match, 1.25%; Figure 3). To our surprise, we found that more than 80% of *Wolbachia*-specific hits were sequences from various *Wolbachia* strains of supergroup B (H03: 772 matches, 83.8% and H05: 65 matches, 81.3%) rather than of supergroup A (H03: 118 matches, 12.8% and H05: 14 matches, 17.5%) where wMel and wMelCS belong to. A small subset of the matches further belonged to supergroup T (Figure 3).

**Figure 3.**
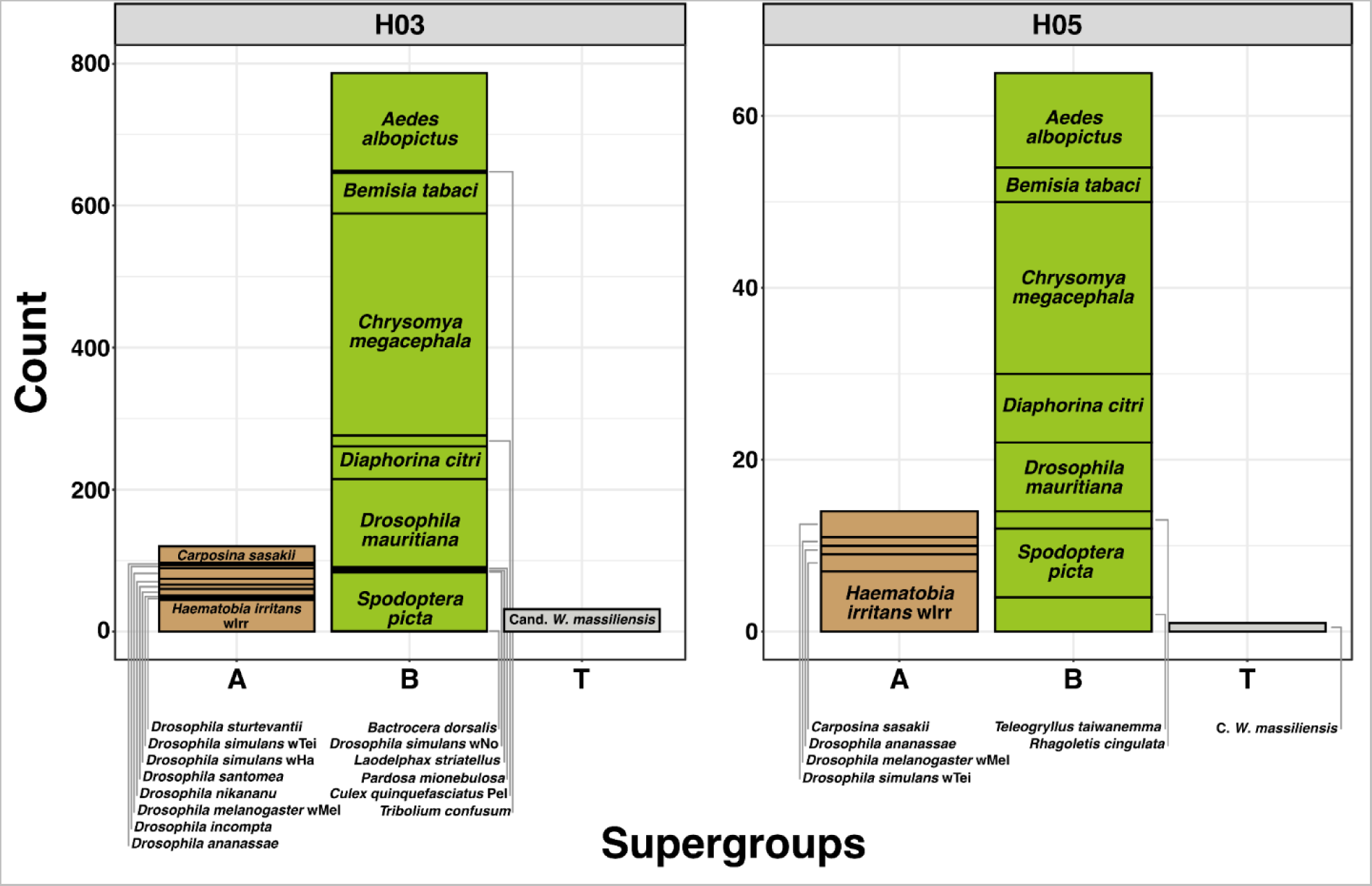
BLAST hits of raw Illumina reads from samples H03 and H05. Stacked barplots showing the number of BLAST hits, with identity values >99% to *Wolbachia*-specific target sequences from different *Wolbachia* types. These belong to supergroups A (brown), B (green) or T (grey). The labels associated with the stacked boxes indicate the names of the host species for each *Wolbachia* type and the size of the stacked boxes indicate the number of reads with matches to the corresponding *Wolbachia* type.

These findings suggest that the two samples H03 and H05 were infected with yet unknown *Wolbachia* types of supergroup B. Consistent with this hypothesis, we indeed found that samples H03 and H05 preferentially mapped to the supergroup B *Wolbachia* type wMeg (Figure 4), when mapping raw reads against a joint reference consisting of wMeg and wMelCS. Mapping statistics revealed that 50% and 24.4% of the wMeg reference genome were covered by an average 8.8-fold and 2.2-fold RD for samples H03 and H05, respectively (Figure 4). Conversely, reference coverages for the five control samples were markedly lower (average 4.4%) and never exceeded 7.65% (for sample H20).

**Figure 4.**
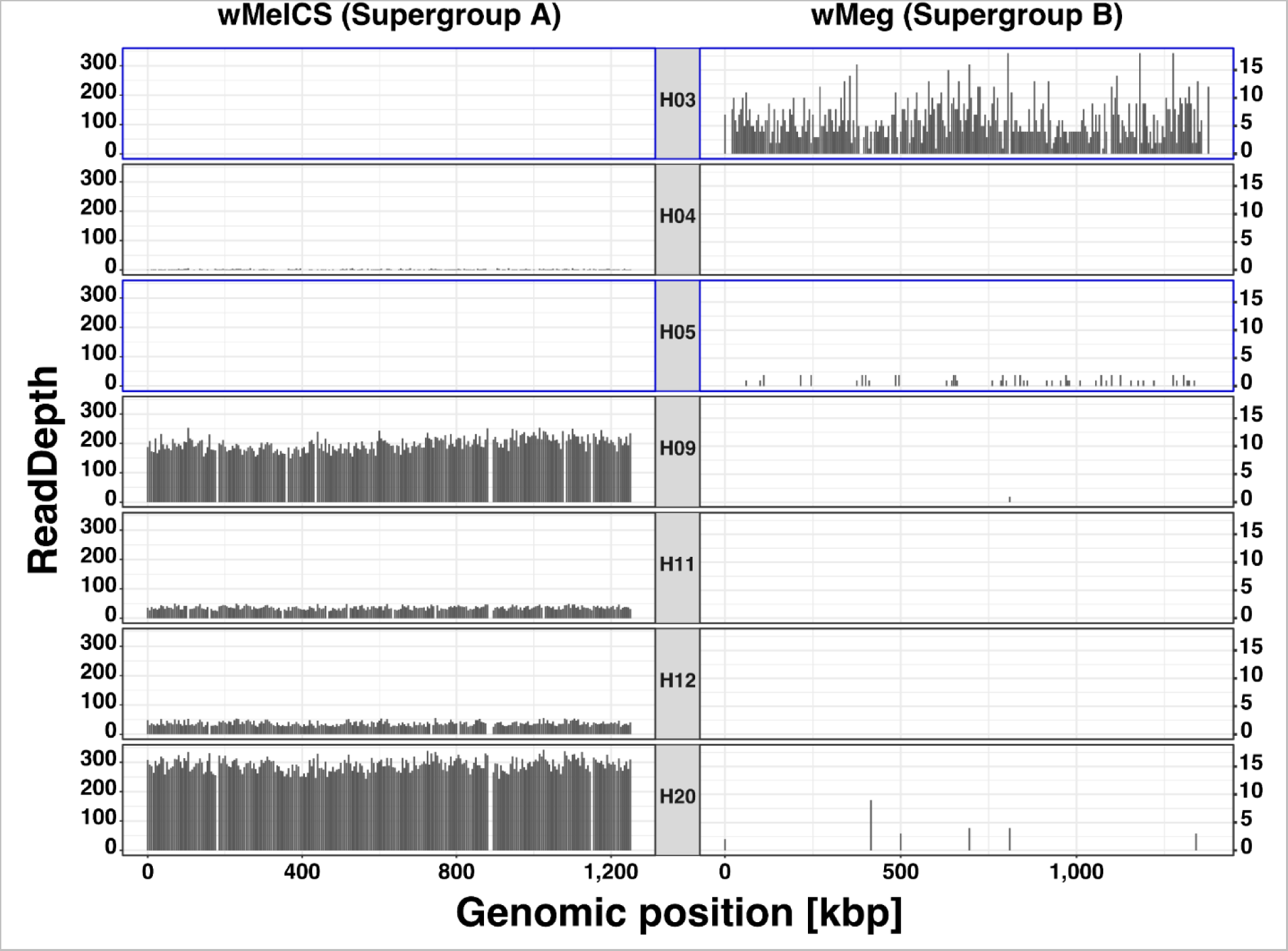
Read depths (RD) along reference genomes from supergroups A and B. Barplots showing median RD in 5,000 bp non-overlapping windows for seven historic samples where reads were mapped against a joint reference consisting of the genomes from wMelCS (supergroup A, left) and wMeg (supergroup B, right). The two focal samples H03 and H05, which were presumably infected with a *Wolbachia* type of supergroup B, are outlined in blue. Note, we removed windows with RD exceeding 3 x standard deviation to improve readability.

In spite of these findings, we cannot fully rule out that the presence of *Wolbachia* types from supergroup B in the sequencing data is the result of cross-contamination. For example, endoparasitic nematodes (Taylor *et al*., 2005), ectoparasitic mites (Breeuwer & Jacobs, 1996) or parasitoid wasps (Duplouy *et al*., 2015) that were caught and preserved together with the museum specimens may have brought along their own *Wolbachia* types, which could be wrongly ascribed to the *Drosophila* sample when the animals were sequenced together. Particularly parasitic wasps, such as *Nasonia vitripennis* have *Wolbachia* types that belong to supergroup B (Tiwary *et al*., 2022). Although very unlikely, pest species in the museum, such as the museum beetle *Anthrenus museorum* may also contaminate specimens with DNA of their own *Wolbachia* types when feeding or breeding on museum samples (Kageyama *et al*., 2010). However, since our BLAST search did not show matches with *Wolbachia* types associated with any of the organism groups mentioned above (Figure 3), our findings are a strong argument that the identified *Wolbachia* types were indeed the endosymbionts of the *D. melanogaster* specimens H03 and H05. While infections with different types of supergroup A and B have been previously reported for *Drosophila simulans* (Poinsot & Merçot, 1997), our results provide the first evidence that also its sister species *D. melanogaster* can be naturally infected with supergroup B types - at least in historic samples, which probably predate the invasion of wMel in natural populations.

### 3.3 Phylogenetic relationships between historic and contemporary Wolbachia

To better understand the evolutionary history of historic and contemporary *Wolbachia* samples, we mapped raw sequencing data prefiltered for *Wolbachia*-specific reads with Kraken against the *Wolbachia* reference genome and used 279 genome-wide SNPs for phylogenetic inference (Figure 5). We only included samples into this analysis, which contained sufficient RD at the majority of the SNPs as described in Materials and Methods. Our phylogenetic analyses, which were based on previously available data (including seven of the historic samples) as well as genomic data newly generated by ONT sequencing in this study (nine samples), identified the same four phylogenetic clusters of wMel types in contemporary samples as previously described in Richardson et al. (2012). This confirms the robustness of our SNP-based phylogenetic approach. Furthermore, and consistent with previous data, we find that wMelCS types form a monophyletic cluster distinct from wMel. Consistent with our characterization based on diagnostic maker SNPs as described above, we found that all seven historic samples included in this analysis clustered with the contemporary wMelCS samples. The sample H9, which was collected in Germany in the middle of the 19th century formed a separate branch within the wMelCS cluster. All other historic samples, which were all collected in 1933 in Sweden were positioned in a well-supported monophyletic cluster, which we now denote as Group VIb, that was distinct to all contemporary wMelCS samples (Group VIa; Figure 5). Consistent with their spatiotemporal relatedness, most samples in this cluster were very similar in terms of genetic variation. In contrast, only the branch lengths of samples H07 and H09 were considerably longer than any other terminal branches. Since these two samples had relatively high RD (>200) compared to the other five samples (see Table 2), we do not assume that the high degree of SNP differences in these samples is likely a technical artifact of the lower sequencing depth resulting in false genotypes but rather a true biological signal or the result of advanced DNA degradation in these two samples.

**Figure 5.**
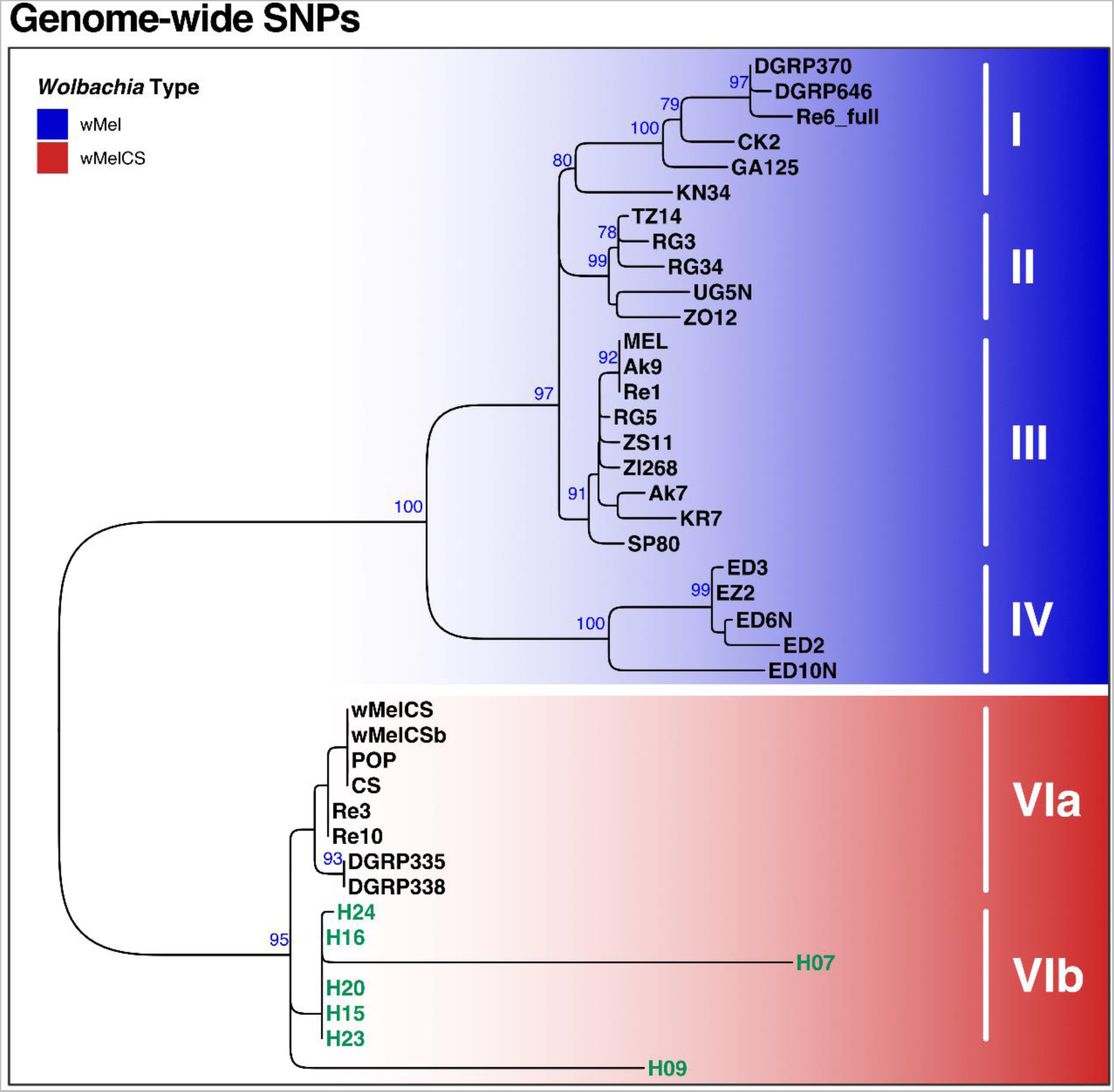
Phylogenetic tree of *Wolbachia* based on 279 genome-wide SNPs. ML tree based on SNP-data from 40 samples of historic and contemporary samples which were either infected with wMel (blue) or wMelCS (red). Raw reads of each sample were mapped against the wMel reference genome (RefSeq: AE017196.1). Roman numbers indicate subgroups of *D. melanogaster*-specific *Wolbachia* strains as previously defined by Richardson et al. (2012). The six historic samples included in this analysis are highlighted in green. Blue numbers indicate bootstrap values >75% from 100 rounds of bootstrapping.

To further investigate the phylogenetic signal in the two samples H03 and H05, which appear to be more closely related to *Wolbachia* types of supergroup B than to wMel and wMelCS of supergroup A, as described above, we repeated SNP calling with relaxed parameters, i.e. we included SNPs with a minimum coverage ≥ 2 and allowed samples with ≥ 80% missing data. As expected from the BLAST results described before, both samples neither clustered with the wMel nor with the wMelCS samples (Figure S1), but rather formed a distinct outgroup. The branch lengths of these samples were extremely long, which indicates substantial sequence divergence to the other wMel and wMelCS types. Consistent with the BLAST results described above, these findings thus further support that samples H03 and H05 were putatively infected with unknown and highly diverged *Wolbachia* types.

### 3.5 Wolbachia and mitochondria of D. melanogaster hosts share a common evolutionary history

Previous analyses have unveiled a close link between the evolutionary histories of *Wolbachia* and mitochondria of the host species (Nunes *et al*., 2008; Richardson *et al*., 2012; Ilinsky, 2013). Both the organelle and the endosymbiont are jointly transmitted maternally which results in a strong influence of *Wolbachia* on the distribution of genetic variation in mitochondria (Richardson *et al*., 2012). This pattern of co-evolution may be only interrupted by sporadic events of horizontal transmission. Here, we tested if we observe similar effects in historic and contemporary *D. melanogaster* samples by applying the same SNP-based phylogenetic analyses to mitochondrium-specific reads as described above. A direct comparison of the topologies from mitochondrial and *Wolbachia* trees as shown in the tanglegram in Figure 6 further reveals the tight links between the evolutionary histories of mitochondria and *Wolbachia*. The only major difference between the two trees is the placement of sample H09, which is placed at the base of the wMelCS cluster in the *Wolbachia* tree but located within the wMelCS cluster in the mitochondrial tree.

**Figure 6.**
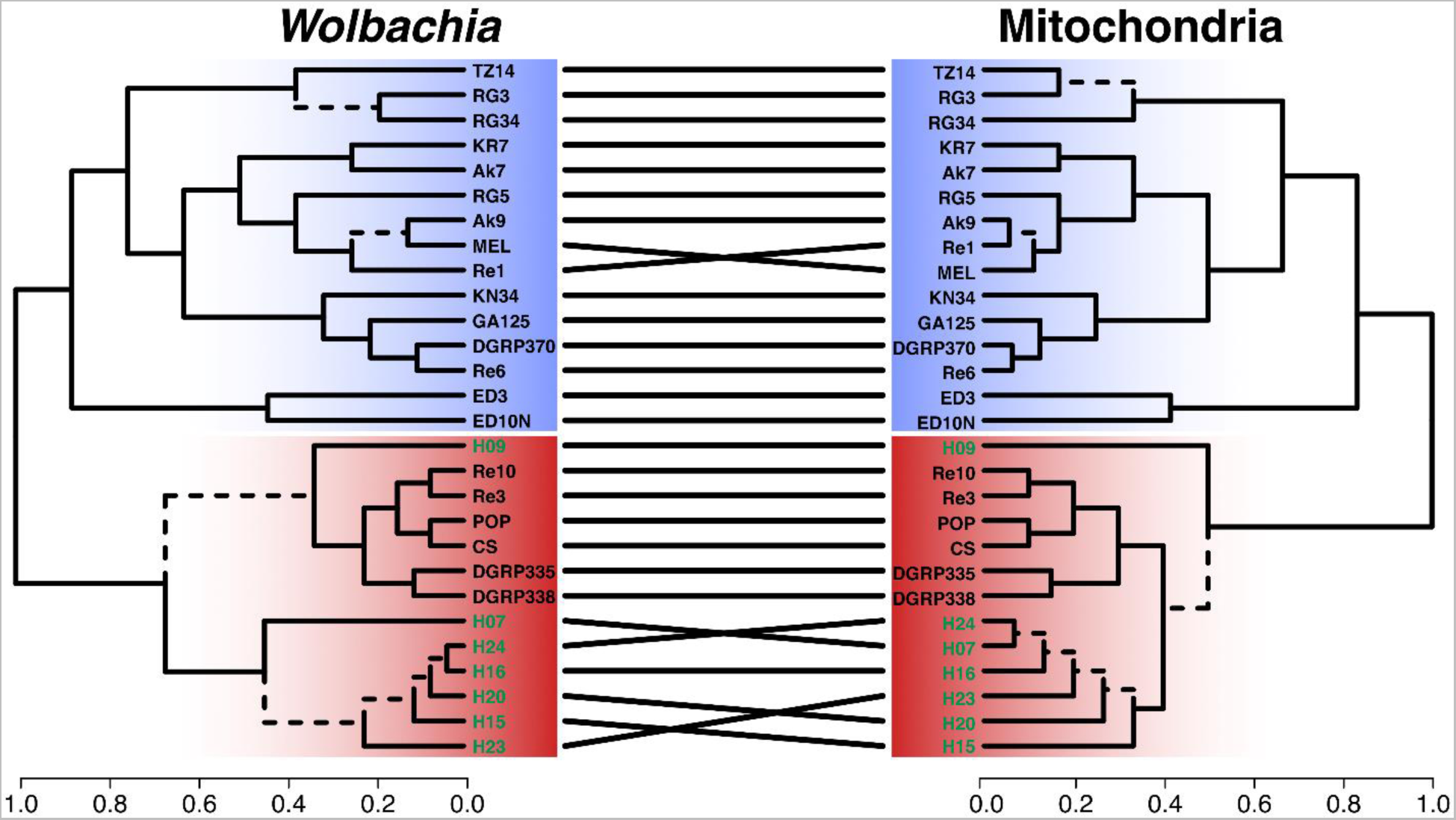
Tanglegram comparing the phylogenetic trees of *Wolbachia* and mitochondria of *D. melanogaster*. Ultrametric trees were generated from SNP-based ML trees for *Wolbachia* and mitochondria of the same historic (highlighted in green) and contemporary samples. Dashed edges indicate topology differences between the trees. Samples of type wMel are highlighted with a blue background and samples of type wMelCS with a red background.

Consistent with our expectations, we found that mitochondrial samples clustered according to their *Wolbachia-*type, i.e., historic and contemporary wMelCS-infected samples formed a monophyletic cluster that was distinct from the contemporary wMel samples (Figure 7). Curiously, we found that even the uninfected historic mitochondrial samples were located within the wMelCS cluster. This provides further evidence that modern mitochondrial haplotypes which are nowadays associated with wMel were at least rather rare in Northern Europe 100-200 years ago. In contrast to all other historic samples and to our surprise, we found only one sample, H05, which was putatively infected with an unknown *Wolbachia* type as described above, clustering basal to all wMel-specific mitochondrial haplotypes. Sample H05 was collected in Denmark more than 200 years ago, which presumably predates the wMel invasion and falls in the same time-period when the first European *Drosophila melanogaste*r just started to appear in North America (Keller, 2007). We therefore speculate that sample H05, in spite of not being infected with wMel, belonged to the same mitochondrial haplogroup from which later the worldwide invasion of wMel started. In contrast to the hypothesis put forward in Riegler et al.(2005), we thus assume that the wMel invasion began in the old world with hosts that carried mitochondrial haplotypes similar to the 200-year-old Danish fly H05, rather than originating from North America.

**Figure 7.**
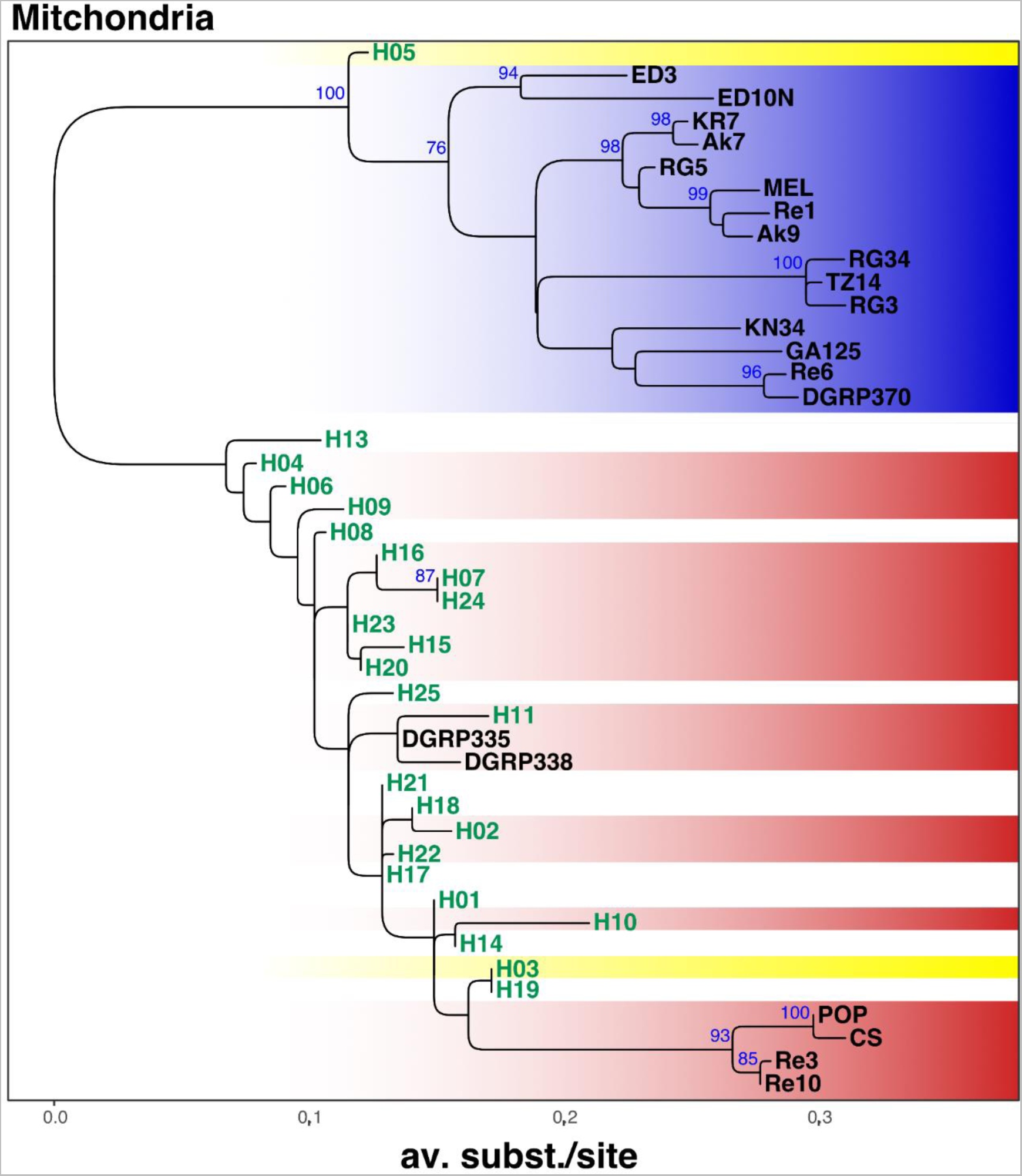
Phylogenetic tree of mitochondria based on 291 genome-wide SNPs. ML tree (mid-point rooted) based on SNP-data from 45 samples of historic and contemporary samples which were either infected with wMel (blue) or wMelCS (red). The two samples H03 and H05 with uncertain *Wolbachia* type are highlighted in yellow and uninfected specimens are shown in white. The 24 historic samples that were included in this analysis are highlighted in green. Blue numbers indicate bootstrap values >75% from 100 rounds of bootstrapping.

### 3.6 De-novo genome assemblies of historic and contemporary Wolbachia genomes

Finally, we further used the sequencing data of the ten historic samples, which we classified as infected, and of the nine newly sequenced contemporary samples for de-novo genome assembly. Assembly quality, which we assessed based on the BUSCO approach and on general assembly statistics, such as total number of contigs, cumulative contig lengths, length of largest contig, N50 and N90, varied highly among the different datasets (Table 2). For five of the historic samples (H15, H16, H20, H23 and H24) we were able to produce de-novo assemblies where the cumulative length of *Wolbachia*-specific contigs were within the range of the expected total genome size (i.e., approximately 1.26 mb). In contrast, cumulative contig lengths for the other samples were markedly shorter ranging from 26 kb to 590 kb in spite of RD > 10-fold. This suggests elevated levels of sequence degradation in these samples which is further indicated by N50 values < 100 bp and GC ratios, which are markedly higher than the *wMel*-specific GC-ratio of 35.2%.

**Table 3.** Summary statistics of de-novo assemblies of *Wolbachia* genomes in historic and recent samples. based on Illumina sequencing (published data, historic samples) and ONT sequencing (this study, recent samples).

| Sample ID | Age | #Contigs | GC [%] | Largest Contig [bp] | Total length [bp] | N50 [bp] | N90 [bp] |
| --- | --- | --- | --- | --- | --- | --- | --- |
| H04 | Historic | 34 | 42.31 | 2,074 | 26,114 | 808 | 536 |
| H07 | Historic | 736 | 35.65 | 3,086 | 598,293 | 807 | 547 |
| H09 | Historic | 888 | 35.09 | 4,474 | 893,957 | 1,098 | 601 |
| H11 | Historic | 170 | 41.22 | 1,689 | 122,320 | 698 | 526 |
| H12 | Historic | 116 | 40.57 | 2,400 | 88,387 | 738 | 536 |
| H15 | Historic | 344 | 36.11 | 23,477 | 1,342,387 | 9,929 | 1,391 |
| H16 | Historic | 434 | 35.97 | 18,847 | 1,320,501 | 6,441 | 1,088 |
| H20 | Historic | 608 | 35.75 | 17,791 | 1,283,117 | 3,321 | 869 |
| H23 | Historic | 328 | 35.94 | 36,386 | 1,352,402 | 14,530 | 1,108 |
| H24 | Historic | 490 | 35.74 | 20,317 | 1,281,146 | 4,904 | 941 |
| Ak7 | Recent | 28 | 35.29 | 186,712 | 1,261,419 | 105,326 | 27,572 |
| Ak9 | Recent | 27 | 35.41 | 209,106 | 1,317,183 | 125,544 | 32,583 |
| CS | Recent | 33 | 35.3 | 226,845 | 1,252,003 | 86,121 | 17,045 |
| MEL | Recent | 34 | 35.29 | 135,716 | 1,284,797 | 86,112 | 19,843 |
| POP | Recent | 29 | 35.3 | 238,089 | 1,266,220 | 124,125 | 30,868 |
| Re1 | Recent | 23 | 35.35 | 319,617 | 1,268,608 | 158,063 | 31,013 |
| Re10 | Recent | 28 | 35.26 | 365,624 | 1,247,994 | 124,743 | 24,669 |
| Re3 | Recent | 22 | 35.28 | 347,181 | 1,260,048 | 116,215 | 37,219 |
| Re9 | Recent | 25 | 35.68 | 301,634 | 1,326,130 | 158,072 | 31,014 |

Consistent with variable data quality of the investigated historic samples, we found large differences in the proportion of complete BUSCO sequences for the different assemblies (Figure 8). The five high-quality assemblies were characterized by >80% complete and <10% missing BUSCO genes. Conversely, BUSCO results for the other five samples ranged from a complete absence of even fragmented BUSCO genes (H04) to <40% complete BUSCO (H09). Pronounced differences in assembly quality are mostly determined by RD, but may also be influenced by differences in the DNA quality, i.e. due to DNA contamination and degradation. This would also be consistent with longer branches of samples H07 and H09 in the SNP-based phylogenetic analysis (Figure 5) which suggest a pronounced number of private SNPs in these two samples.

**Figure 8.**
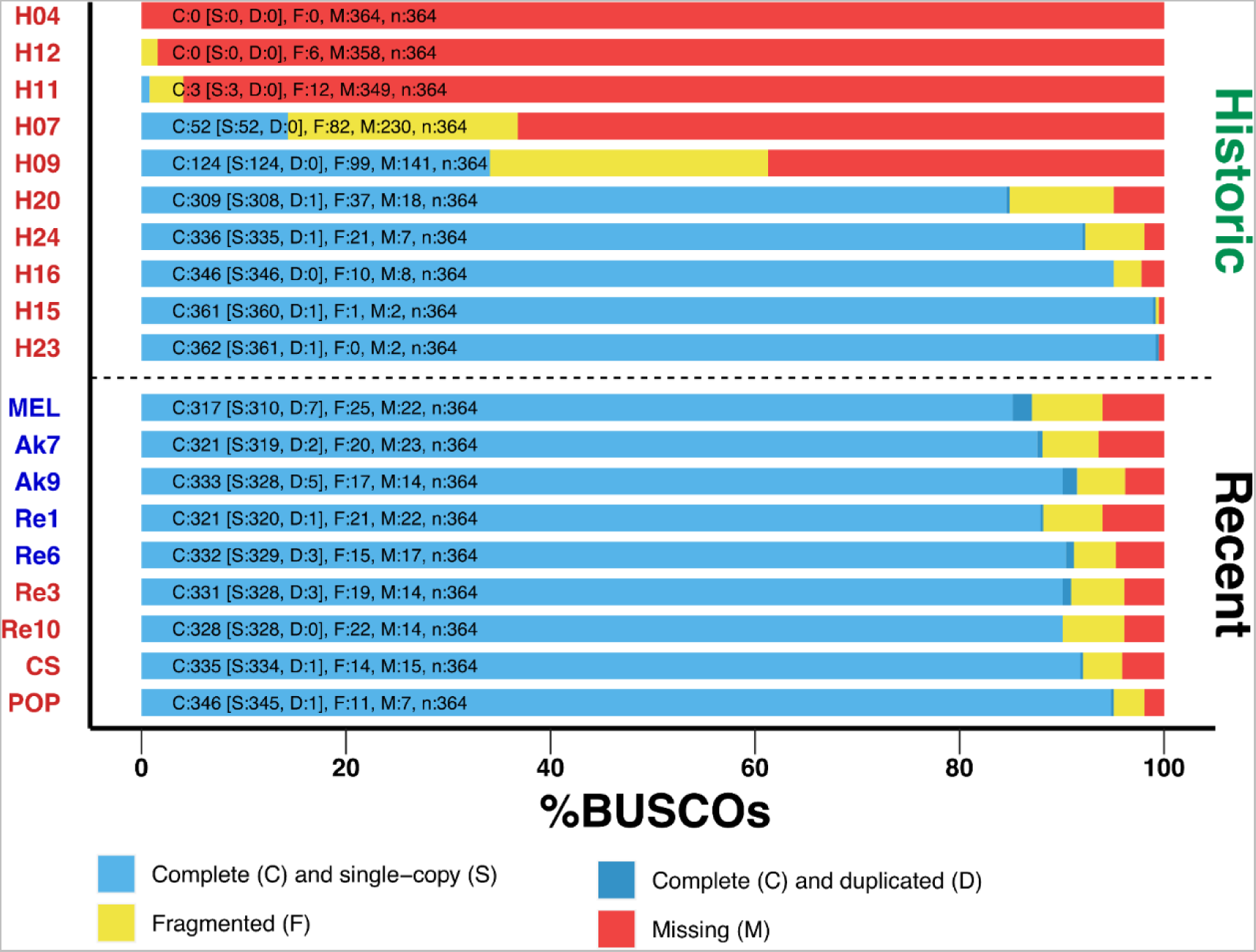
BUSCO scores of de novo assemblies. Stacked barplots showing the proportions of complete, complete and duplicated, fragmented and missing BUSCO genes of the order Rickettsiales identified in each of the assembled genomes. Historic samples are shown above the dashed lines and nine newly sequenced specimens are shown below the line. The color of the label names indicate that they are either of type wMel (blue) or wMelCS (red).

Complementary to the SNP-based phylogenetic analysis, described above, we further aligned 104 BUSCO genes that were complete and present in most of the de-novo assembled genomes of historic samples, the newly sequenced contemporary samples and in the RefSeq datasets included in our analyses. In addition, we included the wYak reference genome as an outgroup and reconstructed a ML tree based on the concatenated gene alignments, which yielded a total length of 68,992bp. The resulting tree, which includes five historic samples that had sufficient RD and data quality to assemble draft genomes, qualitatively confirms our previous findings (Figure 9). Notably, all five historic samples cluster with the contemporary wMelCS samples, which provides further support that they were also of wMelCS type.

**Figure 9.**
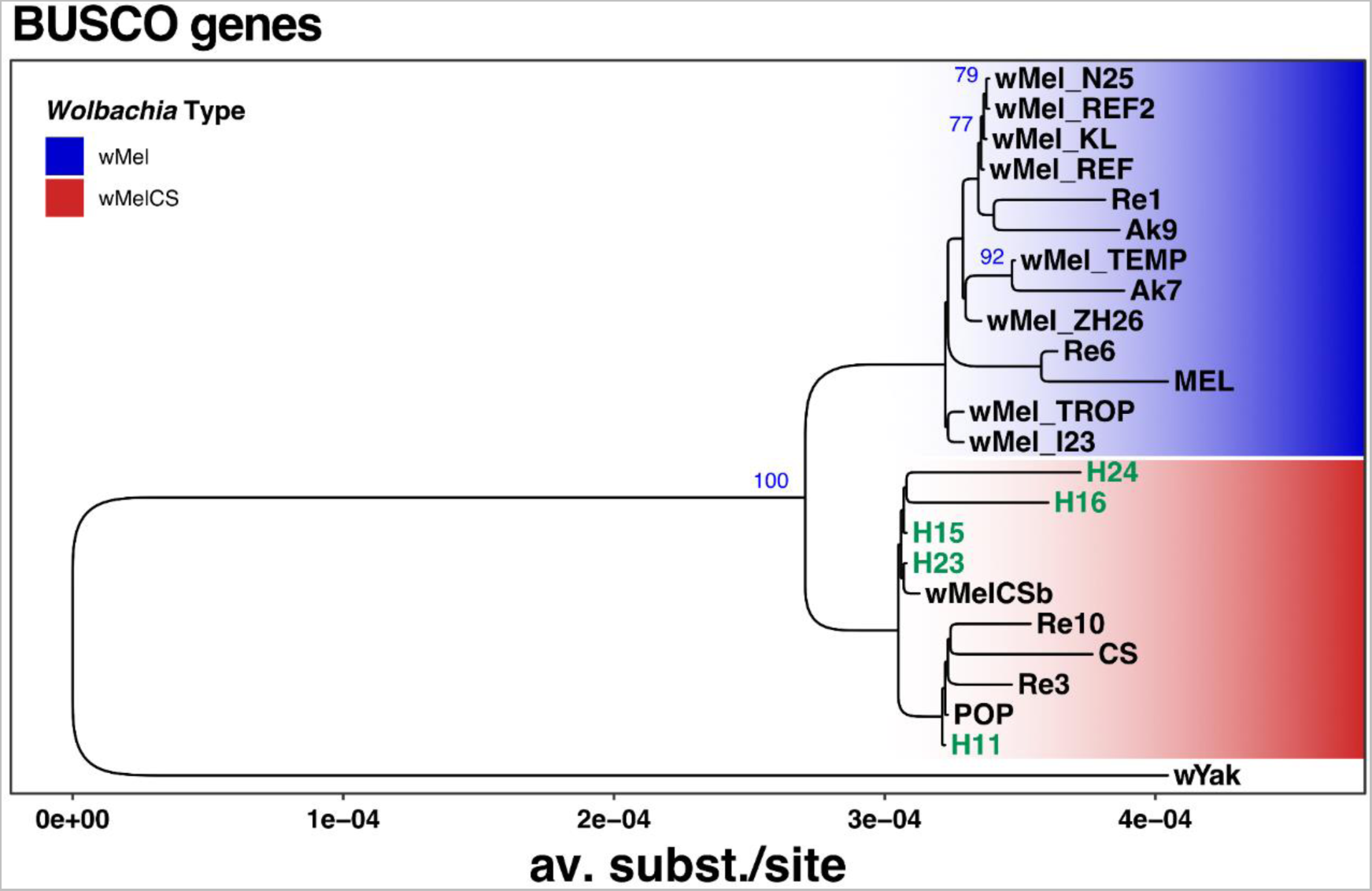
Phylogenetic tree of *Wolbachia* based on 104 aligned and concatenated BUSCO genes. ML tree based on concatenated sequence alignments of 104 genes from 23 historic and contemporary samples which were either infected with wMel (blue) or wMelCS (red) and wYak as an outgroup. The five historic samples that were included in this analysis are highlighted in green. Blue numbers indicate bootstrap values >75% from 100 rounds of bootstrapping.

## 4. Conclusions

In summary, we for the first time identified and characterized historic *Wolbachia* infections in *Drosophila melanogaster* samples that were collected between 90 and more than 200 years ago using publicly available high-quality shotgun sequencing data. Our comparison of historic and contemporary *Wolbachia* samples provides unique empirical evidence that *Wolbachia* infections in Northern European *Drosophila melanogaster* populations were dominated by wMelCS-type bacteria. These findings strongly support the hypothesis put forward in Riegler et al. (2005), that the ancestral wMelCS type was recently replaced by the nowadays common wMel type. While our evidence is only based on eleven samples that were all characterized as wMelCS, these samples were collected across two centuries in Northern Europe and thus span a broad geographic and temporal range. Nevertheless, given that our geographic sampling is limited to few in individuals in Sweden, one sample in Denmark and one sample in Denmark, we cannot rule out that wMel already occurred at lower frequencies in the sampled populations or elsewhere in Europe at the time point they were collected. In addition, we for the first time identify potential historic infections with *Wolbachia* types of supergroup B, which may have important implications on our future understanding of the joint evolutionary history of *Wolbachia* and *Drosophila melanogaster*. Complementary to several recent studies, which investigated *Wolbachia* evolution on a broad taxonomic (Wu *et al*., 2004; Scholz *et al*., 2020) and geographic scale (Richardson *et al*., 2012; Early & Clark, 2013; Basting & Bergman, 2019; Scholz *et al*., 2020), our analyses provide novel insights into the evolutionary history of *Wolbachia* infections in *Drosophila melanogaster* on a broad temporal scale. We are convinced that the unique resource provided by Shpak et al. (2023) represents a seminal data source and an important proof of concept. Their work and spin-off projects like this study or the study by Scarpa et al. (2023), who used this dataset to study the evolutionary history of transposable elements in *D. melanogaster*, will stimulate further quantitative research based on deep sequencing of museum samples (museomics). Such new datasets will enable studies to comprehensively quantify and investigate evolutionary change by comparing historic to contemporary genomic data in model and non-model organisms preserved in museum collections.

## Acknowledgments

We thank Max Shpak, John Pool and Marcus Stensmyr for generously making the genomes of the historical specimens available to the public. We further thank John Pool and Marcus Stensmyr for discussions and their support. We are grateful to Wolfgang Miller, who provided lab space, the laboratory strains used for ONT sequencing and who conceptually supported this project. We thank the DrosEU consortium who provided us with the natural fly lines from Portugal and Finland used for ONT sequencing. We further want to acknowledge Elina Koivisto, Jasmin Jester and Marlene Mühlbauer who helped with fly maintenance and fly food cooking. This project was funded by a standalone grant from the Austrian Science Fund (FWF P32275) to Martin Kapun.

## Authors contributions

Authors contribution according to the CRediT scheme; AS: Conceptualization, Methodology, Validation, Investigation, Writing - Original Draft, SK: Investigation, Writing - Review & Editing, JS: Investigation, Writing - Review & Editing, LK: Writing - Review & Editing, Supervision, Resources, EH: Conceptualization, Writing - Original Draft, Supervision, Resources, MK: Conceptualization, Methodology, Software, Formal Analysis, Resources, Data Curation, Writing - Original Draft, Visualization, Supervision, Project administration, Funding acquisition; The authors declare no competing interests

## Data availability

The full analysis pipeline for this project and the draft genomes can be found at https://github.com/capoony/WolbachiaEvolHist_2023. The newly generated raw ONT sequencing data can be found at the NCBI Short Read Archive (SRA) under the project number: PRJNA987350.

## References

Bandi, C., Damiani, G., Magrassi, L., Grigolo, A., Fani, R. & Sacchi, L. (1994) Flavobacteria as intracellular symbionts in cockroaches. *Proceedings*. Biological Sciences, 257, 43–48.

Bankevich, A., Nurk, S., Antipov, D., Gurevich, A.A., Dvorkin, M., Kulikov, A.S., et al. (2012) SPAdes: A New Genome Assembly Algorithm and Its Applications to Single-Cell Sequencing. Journal of Computational Biology, 19, 455–477.

Basting, P.J. & Bergman, C.M. (2019) Complete Genome Assemblies for Three Variants of the Wolbachia Endosymbiont of Drosophila melanogaster. Microbiology Resource Announcements, 8, e00956–19.

Breeuwer, J.A. & Jacobs, G. (1996) Wolbachia: intracellular manipulators of mite reproduction. Experimental & Applied Acarology, 20, 421–434.

Camacho, C., Coulouris, G., Avagyan, V., Ma, N., Papadopoulos, J., Bealer, K., et al. (2009) BLAST+: architecture and applications. BMC bioinformatics, 10, 421.

Chrostek, E., Marialva, M.S.P., Esteves, S.S., Weinert, L.A., Martinez, J., Jiggins, F.M., et al. (2013) Wolbachia Variants Induce Differential Protection to Viruses in Drosophila melanogaster: A Phenotypic and Phylogenomic Analysis. PLOS Genetics, 9, e1003896.

Chrostek, E., Martins, N., Marialva, M.S. & Teixeira, L. (2021) Wolbachia-Conferred Antiviral Protection Is Determined by Developmental Temperature. mBio, 12, e02923–20.

Correa, C.C. & Ballard, J.W.O. (2016) Wolbachia Associations with Insects: Winning or Losing Against a Master Manipulator. Frontiers in Ecology and Evolution, 3.

Danecek, P., Bonfield, J.K., Liddle, J., Marshall, J., Ohan, V., Pollard, M.O., et al. (2021) Twelve years of SAMtools and BCFtools. GigaScience, 10, giab008.

Dedeine, F., Vavre, F., Fleury, F., Loppin, B., Hochberg, M.E. & Boulétreau, M. (2001) Removing symbiotic Wolbachia bacteria specifically inhibits oogenesis in a parasitic wasp. Proceedings of the National Academy of Sciences, 98, 6247–6252.

Duarte, E.H., Carvalho, A., López-Madrigal, S., Costa, J. & Teixeira, L. (2021) Forward genetics in Wolbachia: Regulation of Wolbachia proliferation by the amplification and deletion of an addictive genomic island. PLoS Genetics, 17, e1009612.

Duplouy, A., Couchoux, C., Hanski, I. & Nouhuys, S. van. (2015) Wolbachia Infection in a Natural Parasitoid Wasp Population. PLOS ONE, 10, e0134843.

Early, A.M. & Clark, A.G. (2013) Monophyly of Wolbachia pipientis genomes within Drosophila melanogaster: geographic structuring, titre variation and host effects across five populations. Molecular Ecology, 22, 5765–5778.

Galili, T. (2015) dendextend: an R package for visualizing, adjusting and comparing trees of hierarchical clustering. Bioinformatics, 31, 3718–3720.

Gerth, M., Gansauge, M.-T., Weigert, A. & Bleidorn, C. (2014) Phylogenomic analyses uncover origin and spread of the Wolbachia pandemic. Nature Communications, 5, 5117.

Gomes, T.M.F.F., Wallau, G.L. & Loreto, E.L.S. (2022) Multiple long-range host shifts of major Wolbachia supergroups infecting arthropods. Scientific Reports, 12, 8131.

Hedges, L.M., Brownlie, J.C., O’Neill, S.L. & Johnson, K.N. (2008) Wolbachia and Virus Protection in Insects. Science, 322, 702–702.

Hertig, M. (1936) The Rickettsia, Wolbachia pipientis (gen. et sp.n.) and Associated Inclusions of the Mosquito, Culex pipiens. Parasitology, 28, 453–486.

Hoffmann, A.A., Turelli, M. & Simmons, G.M. (1986) Unidirectional incompatibility between populations of Drosophila simulans. Evolution, 40, 692–701.

Hoskins, R.A., Carlson, J.W., Wan, K.H., Park, S., Mendez, I., Galle, S.E., et al. (2015) The Release 6 reference sequence of the Drosophila melanogaster genome. Genome Research, 25, 445–458.

Hosokawa, T., Koga, R., Kikuchi, Y., Meng, X.-Y. & Fukatsu, T. (2010) Wolbachia as a bacteriocyte-associated nutritional mutualist. Proceedings of the National Academy of Sciences, 107, 769–774.

Ilinsky, Y. (2013) Coevolution of Drosophila melanogaster mtDNA and Wolbachia Genotypes. PLoS ONE, 8, e54373.

Kageyama, D., Narita, S., Imamura, T. & Miyanoshita, A. (2010) Detection and identification of Wolbachia endosymbionts from laboratory stocks of stored-product insect pests and their parasitoids. Journal of Stored Products Research, 46, 13–19.

Kapun, M., Barrón, M.G., Staubach, F., Obbard, D.J., Wiberg, R.A.W., Vieira, J., et al. (2020) Genomic Analysis of European Drosophila melanogaster Populations Reveals Longitudinal Structure, Continent-Wide Selection, and Previously Unknown DNA Viruses. Molecular Biology and Evolution, 37, 2661–2678.

Katoh, K. & Standley, D.M. (2013) MAFFT Multiple Sequence Alignment Software Version 7: Improvements in Performance and Usability. Molecular Biology and Evolution, 30, 772–780.

Kaur, R., Shropshire, J.D., Cross, K.L., Leigh, B., Mansueto, A.J., Stewart, V., et al. (2021) Living in the endosymbiotic world of Wolbachia: A centennial review. Cell Host & Microbe.

Keller, A. (2007) Drosophila melanogaster’s history as a human commensal. Current Biology, 17, R77–R81.

Kolmogorov, M., Yuan, J., Lin, Y. & Pevzner, P.A. (2019) Assembly of long, error-prone reads using repeat graphs. Nature Biotechnology, 37, 540–546.

Kozlov, A.M., Darriba, D., Flouri, T., Morel, B. & Stamatakis, A. (2019) RAxML-NG: a fast, scalable and user-friendly tool for maximum likelihood phylogenetic inference. Bioinformatics, 35, 4453–4455.

Lack, J.B., Cardeno, C.M., Crepeau, M.W., Taylor, W., Corbett-Detig, R.B., Stevens, K.A., et al. (2015) The *Drosophila* Genome Nexus: A Population Genomic Resource of 623 *Drosophila melanogaster* Genomes, Including 197 from a Single Ancestral Range Population. Genetics, 199, 1229–1241.

Lack, J.B., Lange, J.D., Tang, A.D., B, C.-D., Russell & Pool, J.E. (2016) A Thousand Fly Genomes: An Expanded Drosophila Genome Nexus. Molecular Biology and Evolution, 33, msw195–3313.

Laetsch, D.R. & Blaxter, M.L. (2017) BlobTools: Interrogation of genome assemblies.

Landmann, F. (2019) The Wolbachia Endosymbionts. Microbiology Spectrum, 7.

Lefoulon, E., Bain, O., Makepeace, B.L., Haese, C. d’, Uni, S., Martin, C., et al. (2016) Breakdown of coevolution between symbiotic bacteria Wolbachia and their filarial hosts. PeerJ, 4, e1840.

Li, H. (2013) Aligning sequence reads, clone sequences and assembly contigs with BWA-MEM. *arXiv*:1303.3997 [q-bio].

Li, H. (2018) Minimap2: pairwise alignment for nucleotide sequences. Bioinformatics, 34, 3094– 3100.

Li, H., Handsaker, B., Wysoker, A., Fennell, T., Ruan, J., Homer, N., et al. (2009) The Sequence Alignment/Map format and SAMtools. Bioinformatics, 25, 2078–2079.

Lin, Y., Yuan, J., Kolmogorov, M., Shen, M.W., Chaisson, M. & Pevzner, P.A. (2016) Assembly of long error-prone reads using de Bruijn graphs. Proceedings of the National Academy of Sciences, 113, E8396–E8405.

Lindsey, A.R.I., Bordenstein, S.R., Newton, I.L.G. & Rasgon, J.L. (2016) Wolbachia pipientis should not be split into multiple species: A response to Ramírez-Puebla et al., “Species in Wolbachia? Proposal for the designation of ‘Candidatus Wolbachia bourtzisii’, ‘Candidatus Wolbachia onchocercicola’, ‘Candidatus Wolbachia blaxteri’, ‘Candidatus Wolbachia brugii’, ‘Candidatus Wolbachia taylori’, ‘Candidatus Wolbachia collembolicola’ and ‘Candidatus Wolbachia multihospitum’ for the different species within Wolbachia supergroups.” Systematic and applied microbiology, 39, 220–222.

Lo, N., Casiraghi, M., Salati, E., Bazzocchi, C. & Bandi, C. (2002) How Many Wolbachia Supergroups Exist? Molecular Biology and Evolution, 19, 341–346.

Mackay, T.F.C., Richards, S., Stone, E.A., Barbadilla, A., Ayroles, J.F., Zhu, D., et al. (2012) The Drosophila melanogaster Genetic Reference Panel. Nature, 482, 173–178.

Manni, M., Berkeley, M.R., Seppey, M. & Zdobnov, E.M. (2021) BUSCO: Assessing Genomic Data Quality and Beyond. Current Protocols, 1, e323.

Marçais, G., Delcher, A.L., Phillippy, A.M., Coston, R., Salzberg, S.L. & Zimin, A. (2018) MUMmer4: A fast and versatile genome alignment system. PLOS Computational Biology, 14, e1005944.

Martin, M. (2011) Cutadapt removes adapter sequences from high-throughput sequencing reads. EMBnet.journal, 17, 10–12.

Meany, M.K., Conner, W.R., Richter, S.V., Bailey, J.A., Turelli, M. & Cooper, B.S. (2019) Loss of cytoplasmic incompatibility and minimal fecundity effects explain relatively low Wolbachia frequencies in Drosophila mauritiana. Evolution, 73, 1278–1295.

Mikheenko, A., Prjibelski, A., Saveliev, V., Antipov, D. & Gurevich, A. (2018) Versatile genome assembly evaluation with QUAST-LG. *Bioinformatics (Oxford*, England), 34, i142–i150.

Miller, W.J., Ehrman, L. & Schneider, D. (2010) Infectious Speciation Revisited: Impact of Symbiont-Depletion on Female Fitness and Mating Behavior of Drosophila paulistorum. PLoS Pathogens, 6, e1001214.

Min, K.-T. & Benzer, S. (1997) Wolbachia, normally a symbiont of Drosophila, can be virulent, causing degeneration and early death. Proceedings of the National Academy of Sciences, 94, 10792–10796.

Moreira, L.A., Iturbe-Ormaetxe, I., Jeffery, J.A., Lu, G., Pyke, A.T., Hedges, L.M., et al. (2009) A Wolbachia symbiont in Aedes aegypti limits infection with dengue, Chikungunya, and Plasmodium. Cell, 139, 1268–1278.

Neupane, S., Bonilla, S.I., Manalo, A.M. & Pelz-Stelinski, K.S. (2022) Complete de novo assembly of Wolbachia endosymbiont of Diaphorina citri Kuwayama (Hemiptera: Liviidae) using long-read genome sequencing. Scientific Reports, 12, 125.

Nunes, M.D.S., Nolte, V. & Schlötterer, C. (2008) Nonrandom Wolbachia Infection Status of Drosophila melanogaster Strains with Different mtDNA Haplotypes. Molecular Biology and Evolution, 25, 2493–2498.

Olanratmanee, P., Baimai, V., Ahantarig, A. & Trinachartvanit, W. (2021) Novel Supergroup U Wolbachia in bat mites of Thailand. The Southeast Asian Journal of Tropical Medicine and Public Health, 52, 48–55.

O’Neill, S.L. & Karr, T.L. (1990) Bidirectional incompatibility between conspecific populations of Drosophila simulans. Nature, 348, 178–180.

Osborne, S.E., Leong, Y.S., O’Neill, S.L. & Johnson, K.N. (2009) Variation in Antiviral Protection Mediated by Different Wolbachia Strains in Drosophila simulans. PLOS Pathogens, 5, e1000656.

Pietri, J.E., DeBruhl, H. & Sullivan, W. (2016) The rich somatic life of Wolbachia. MicrobiologyOpen, 5, 923–936.

Poinsot, D. & Merçot, H. (1997) Wolbachia Infection in Drosophila simulans: Does the Female Host Bear a Physiological Cost? Evolution, 51, 180–186.

Pool, J.E., B, C.-D., Russell, Sugino, R.P., Stevens, K.A., Cardeno, C.M., Crepeau, M.W., et al. (2012) Population Genomics of Sub-Saharan Drosophila melanogaster: African Diversity and Non-African Admixture. PLoS Genetics, 8, e1003080.

Prjibelski, A., Antipov, D., Meleshko, D., Lapidus, A. & Korobeynikov, A. (2020) Using SPAdes De Novo Assembler. Current Protocols in Bioinformatics, 70.

R Core Team. (2019) R Foundation for Statistical Computing.

Ramírez-Puebla, S.T., Servín-Garcidueñas, L.E., Ormeño-Orrillo, E., Vera-Ponce De León, A., Rosenblueth, M., Delaye, L., et al. (2015) Species in Wolbachia? Proposal for the designation of ‘Candidatus Wolbachia bourtzisii’, ‘Candidatus Wolbachia onchocercicola’, ‘Candidatus Wolbachia blaxteri’, ‘Candidatus Wolbachia brugii’, ‘Candidatus Wolbachia taylori’, ‘Candidatus Wolbachia collembolicola’ and ‘Candidatus Wolbachia multihospitum’ for the different species within Wolbachia supergroups. Systematic and Applied Microbiology, 38, 390–399.

Raxworthy, C.J. & Smith, B.T. (2021) Mining museums for historical DNA: advances and challenges in museomics. Trends in Ecology & Evolution, 36, 1049–1060.

Richardson, M.F., Weinert, L.A., Welch, J.J., Linheiro, R.S., Magwire, M.M., Jiggins, F.M., et al. (2012) Population Genomics of the Wolbachia Endosymbiont in Drosophila melanogaster. PLoS Genetics, 8, e1003129.

Riegler, M., Iturbe-Ormaetxe, I., Woolfit, M., Miller, W.J. & O’Neill, S.L. (2012) Tandem repeat markers as novel diagnostic tools for high resolution fingerprinting of Wolbachia. BMC Microbiology, 12, S12.

Riegler, M., Sidhu, M., Miller, W.J. & O’Neill, S.L. (2005) Evidence for a Global Wolbachia Replacement in Drosophila melanogaster. Current Biology, 15, 1428–1433.

Robinson, J.T., Thorvaldsdóttir, H., Winckler, W., Guttman, M., Lander, E.S., Getz, G., et al. (2011) Integrative genomics viewer. Nature Biotechnology, 29, 24–26.

Rousset, F. & Solignac, M. (1995) Evolution of single and double Wolbachia symbioses during speciation in the Drosophila simulans complex. Proceedings of the National Academy of Sciences of the United States of America, 92, 6389–6393.

Saha, S., Hunter, W.B., Reese, J., Morgan, J.K., Marutani-Hert, M., Huang, H., et al. (2012) Survey of Endosymbionts in the Diaphorina citri Metagenome and Assembly of a Wolbachia wDi Draft Genome. PLOS ONE, 7, e50067.

Sazama, E.J., Bosch, M.J., Shouldis, C.S., Ouellette, S.P. & Wesner, J.S. (2017) Incidence of Wolbachia in aquatic insects. Ecology and Evolution, 7, 1165–1169.

Scarpa, A., Pianezza, R., Wierzbicki, F. & Kofler, R. (2023) Genomes of historical specimens reveal multiple invasions of LTR retrotransposons in Drosophila melanogaster populations during the 19th century.

Scholz, M., Albanese, D., Tuohy, K., Donati, C., Segata, N. & Rota-Stabelli, O. (2020) Large scale genome reconstructions illuminate Wolbachia evolution. Nature Communications, 11, 5235.

Seppey, M., Manni, M. & Zdobnov, E.M. (2019) BUSCO: Assessing Genome Assembly and Annotation Completeness. Gene Prediction, 227–245.

Serga, S., Maistrenko, O., Rozhok, A., Mousseau, T. & Kozeretska, I. (2014) Fecundity as one of possible factors contributing to the dominance of the wMel genotype of Wolbachia in natural populations of Drosophila melanogaster. Symbiosis, 63, 11–17.

Serga, S.V., Maistrenko, O.M., Matiytsiv, N.P., Vaiserman, A.M. & Kozeretska, I.A. (2021) Effects of Wolbachia infection on fitness-related traits in Drosophila melanogaster. Symbiosis, 83, 163–172.

Shpak, M., Ghanavi, H.R., Lange, J.D., Pool, J.E. & Stensmyr, M.C. (2023) Genomes from 25 historical Drosophila melanogaster specimens illuminate adaptive and demographic changes across more than 200 years of evolution.

Singhal, K. & Mohanty, S. (2018) Comparative genomics reveals the presence of putative toxin-antitoxin system in Wolbachia genomes. Molecular genetics and genomics: MGG, 293, 525– 540.

Strunov, A., Lerch, S., Blanckenhorn, W.U., Miller, W.J. & Kapun, M. (2022) Complex effects of environment and Wolbachia infections on the life history of Drosophila melanogaster hosts. Journal of Evolutionary Biology, 35, 788–802.

Taylor, M.J., Bandi, C. & Hoerauf, A. (2005) Wolbachia bacterial endosymbionts of filarial nematodes. Advances in Parasitology, 60, 245–284.

Teixeira, L., Ferreira, Á. & Ashburner, M. (2008) The Bacterial Symbiont Wolbachia Induces Resistance to RNA Viral Infections in Drosophila melanogaster. PLoS Biology, 6, e1000002.

Tiwary, A., Babu, R., Sen, R. & Raychoudhury, R. (2022) Bacterial supergroup-specific “cost” of Wolbachia infections in Nasonia vitripennis. Ecology and Evolution, 12, e9219.

Truitt, A.M., Kapun, M., Kaur, R. & Miller, W.J. (2019) *Wolbachia* modifies thermal preference in *Drosophila melanogaster*. Environmental Microbiology, 21, 3259–3268.

Vaser, R., Sović, I., Nagarajan, N. & Šikić, M. (2017) Fast and accurate de novo genome assembly from long uncorrected reads. Genome Research, 27, 737–746.

Weinert, L.A., Araujo-Jnr, E.V., Ahmed, M.Z. & Welch, J.J. (2015) The incidence of bacterial endosymbionts in terrestrial arthropods. Proceedings of the Royal Society B: Biological Sciences, 282, 20150249.

Werren, J.H., Baldo, L. & Clark, M.E. (2008) Wolbachia: master manipulators of invertebrate biology. Nature Reviews Microbiology, 6, 741–751.

Werren, J.H., Zhang, W. & Guo, L.R. (1995) Evolution and phylogeny of Wolbachia: reproductive parasites of arthropods. *Proceedings*. Biological Sciences, 261, 55–63.

Wick, R.R., Judd, L.M. & Holt, K.E. (2019) Performance of neural network basecalling tools for Oxford Nanopore sequencing. Genome Biology, 20, 129.

Wilkinson, S.P. & Davy, S.K. (2018) phylogram: an R package for phylogenetic analysis with nested lists. Journal of Open Source Software, 3, 790.

Woolfit, M., Iturbe-Ormaetxe, I., Brownlie, J.C., Walker, T., Riegler, M., Seleznev, A., et al. (2013) Genomic Evolution of the Pathogenic Wolbachia Strain, wMelPop. Genome Biology and Evolution, 5, 2189–2204.

Wu, M., Sun, L.V., Vamathevan, J., Riegler, M., Deboy, R., Brownlie, J.C., et al. (2004) Phylogenomics of the Reproductive Parasite Wolbachia pipientis wMel: A Streamlined Genome Overrun by Mobile Genetic Elements. PLoS Biology, 2, e69.

Yu, G., Smith, D.K., Zhu, H., Guan, Y. & Lam, T.T.-Y. (2017) ggtree: an r package for visualization and annotation of phylogenetic trees with their covariates and other associated data. Methods in Ecology and Evolution, 8, 28–36.

Zimmermann, J., Hajibabaei, M., Blackburn, D.C., Hanken, J., Cantin, E., Posfai, J., et al. (2008) DNA damage in preserved specimens and tissue samples: a molecular assessment. Frontiers in Zoology, 5, 18.

Zug, R. & Hammerstein, P. (2012) Still a host of hosts for Wolbachia: analysis of recent data suggests that 40% of terrestrial arthropod species are infected. PloS One, 7, e38544.

Zug, R. & Hammerstein, P. (2015) Bad guys turned nice? A critical assessment of Wolbachia mutualisms in arthropod hosts. Biological Reviews, 90, 89–111.

