## Supplementary figures and images for "Historic museum samples provide evidence for a recent replacement of *Wolbachia* types in European *Drosophila melanogaster*"

### Figure S1

Wolbachia

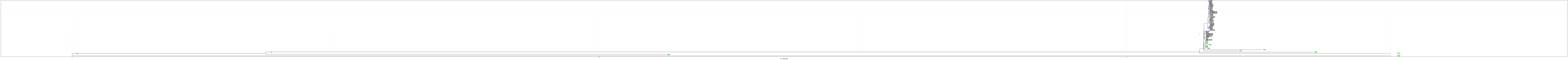
